# Virulence and antimicrobial resistance profile of non-typhoidal *Salmonella enterica* serovars recovered from poultry processing environments at wet markets in Dhaka, Bangladesh

**DOI:** 10.1101/2021.07.23.453547

**Authors:** Nure Alam Siddiky, Md Samun Sarker, Md. Shahidur Rahman Khan, Md. Tanvir Rahman, Md. Abdul Kafi, Mohammed A. Samad

## Abstract

The rapid emergence of virulent and multidrug-resistant (MDR) non-typhoidal *Salmonella* (NTS) *enterica* serovars are a growing public health concern globally. The present study focused on the assessment of the pathogenicity and antimicrobial resistance (AMR) profiling of NTS *enterica* serovars isolated from chicken processing environments at wet markets in Dhaka, Bangladesh. A total number of 870 samples consisting of carcass dressing water (CDW), chopping board swabs (CBS), and knife swabs (KS) were collected from 29 wet markets. The prevalence of *Salmonella* was found to be 20% in CDW, 19.31% in CBS and 17.58% in KS, respectively. Meanwhile, the MDR *Salmonella* was found to be 72.41%, 73.21% and 68.62% in CDW, CBS, and KS, respectively. All isolates were screened by polymerase chain reaction (PCR) for eight virulence genes, namely *inv*A, *agf*A, *Ipf*A, *hil*A, *siv*H, *sef*A, *sop*E, and *spv*C. The *S*. Enteritidis and untyped *Salmonella* isolate harbored all virulence genes while *S*. Typhimurium isolates carried six virulence genes except *sef*A and *spv*C. Phenotypic resistance revealed decreased susceptibility to ciprofloxacin, streptomycin, ampicillin, tetracycline, gentamycin, sulfamethoxazole-trimethoprim, amoxicillin-clavulanic acid and azithromycin. Genotypic resistance showed higher prevalence of plasmid mediated *bla*TEM followed by *tet*A, *sul*1, *sul*2, *sul*3, and *str*A/B genes. Harmonic and symmetrical trend was observed among the phenotypic and genotypic resistance patterns of the isolates. The research findings anticipate that MDR and virulent NTS *enterica* serovars are prevailing in the wet market environments which can easily enter into the human food chain. There was a resilient and significant correlation existent among the phenotypic and genotypic resistance patterns and virulence genes of *Salmonella* isolate recovered from carcass dressing water, chopping board swabs, and knife swabs (*p* < 0.05), respectively.

## Introduction

*Salmonella* has been recognized as one of the common pathogens that causes gastroenteritis (Hawker et al., 2019; Jain et al., 2020) with significant morbidity, mortality, and economic loss (Lin et al. 2014; Sallam et al., 2014). WHO reported 153 million cases of NTS enteric infections worldwide in 2010, of which 56,969 were dead along with 50% were foodborne (Kirk et al., 2015). The disease surveillance report of China from 2006 to 2010 identified *Salmonella* as the second foodborne outbreak (Pang et al., 2011). NTS serovars like Typhimurium and Enteritidis are the predominant worldwide among the 2,600 serotypes of *Salmonella* have been identified (Issenhuth-Jeanjean et al., 2014; Takaya et al., 2019). Poultry has been regarded as the single prime cause of human salmonellosis and avian salmonellosis is not only affects the poultry industry but also can infect humans and caused by the consumption of contaminated poultry meat and eggs (Behravesh et al., 2014). Poultry and eggs are considered to be the primary cause of salmonellosis and numerous other foodborne outbreaks (Gieraltowski et al., 2016; Keerthirathne et al., 2017; Biswas et al., 2019, 2020; Yu et al., 2020). Generally, *Salmonella* grows in animal farms may contaminate eggs and/or meat during the slaughtering process before being transferred to humans through the food chain. Indeed, numerous previous studies have been reported the isolation of *Salmonella* from foods of animal origin as well as human samples (Ed-Dra et al., 2018; Paudyal et al., 2020; Jiang et al., 2019; Elbediwi et al., 2020). Human *S*. Enteritidis are generally linked with the consumption of contaminated eggs and poultry meat, while *S*. Typhimurium with the consumption of pork, poultry, and beef (Park et al., 2014; Spector et al., 2012). Different prevalence of *Salmonella enterica* serovars has been reported around the globe from animal products and by-products (Park et al., 2014; De Freitas et al., 2010; Shah et al., 2012). *Salmonella* Typhimurium and Enteritidis are the most frequently reported serovars associated with human foodborne illnesses (Suresh et al., 2006). Untyped *Salmonella* of animal origin has been increasingly observed in Bangladesh (Momtaz et al., 2018; Sultana et al., 2014) but limited information has been published on *Salmonella enterica* serovars isolated from chicken processing environments and beyond.

Widespread uses of antimicrobials in poultry farming generate benefits for producers but aggravate the emergence of AMR bacteria (Manyi-Loh et al., 2018). Microorganisms that develop AMR are sometimes referred to as superbugs and open the door to treatment failure for even the most common pathogens, raise health care cost, and increases the severity and duration of infections. AMR burden may kill 300 million people during the next 35 years with a terrible impact on the global economy declining GDP by 2-3% in 2050 (O’Neill, 2014). WHO recognized AMR as a serious threat, is no longer a forecast for the future, which is happening around the world and affects everybody regardless of age, sex, and nation (Robicsek et al., 2006). Misuse and overuse of existing antimicrobials in humans, animals, and plants are accelerating the development and spread of AMR (IACG, 2019). The antimicrobials are used in Bangladesh as the therapeutic, preventive and growth promoters in the poultry production system (Al Masud et al., 2020). The problem of AMR *Salmonella* emerged global concern in the modern decade (O’Bryan et al., 2018; Sultana et al., 2014). MDR *Salmonella* of poultry origin have been increasing in Bangladesh (Alam et al., 2020; Siddiky et al., 2021).

Usually, the virulence factors promote the pathogenicity of *Salmonella* infection. Chromosomal and plasmid-mediated virulence factors are associated with the pathogenicity of *Salmonella*. *Salmonella* possesses major virulence genes such as *inv*A, *agf*A, *Ipf*A, *hil*A, *siv*H, *sef*A and *sop*E. The infectivity of *Salmonella* strains is associated with various virulence genes existent in the chromosomal *Salmonella* pathogenicity islands (SPIs) (Nayak et al., 2004). The invasion genes *inv*A, *hil*A, and *siv*H code with a protein in the inner chromosomal membrane of *Salmonella* that is necessary for the invasion to epithelial cells (Darwin et al., 1999). Moreover, *Salmonella* effector protein adhered by *sop*E gene which have relevance to *Salmonella* virulence (Huehn et al., 2010). The plasmid mediated *spv*C gene is responsible for vertical transmission (Silva et al., 2017). The long polar fimbria (*Ipf* operon) make the attraction of the microbes for Peyer’s patches and adhesion to intestinal M cells (Bäumler et al., 1996). The aggregative fimbria (*agf* operon) promote the primary interaction of the *Salmonella* with the intestine of the host and stimulate microbial self-aggregation for higher rates of survival (Collinson et al., 1993). The *Salmonella*-encoded fimbria (*sef* operon) endorses interaction between the microbes and the macrophages (Collinson et al., 1993). Though there was a paucity of information in the determination of virulence gene from *Salmonella enterica* serovars in Bangladesh but recently eight virulence genes were found in *Salmonella* isolates of poultry origin in Bangladesh (Siddiky et al., 2021).

Wet markets are very common in Bangladesh which are commonly dirty, chaotic, and unhygienic and floors are constantly sprayed with water for washing and to conserve the humidity (Nidaullah et al., 2017). Dressing and processing of poultry in the open environment are common practices in the traditional wet markets. The chicken vendors himself dressing the chicken without having personal protective devices, without using clean dressing utensils such as chopping boards and knives. Even same water is used frequently for washing or cleaning the whole dressed carcass. There is a huge chance of cross-contamination and horizontal spreading of MDR *Salmonella* in the wet markets environment (Bupasha et al., 2020). The whole chicken carcass, vendor, and consumer may be infected with *Salmonella* due to poor sanitary and hygienic practices. Even there is a great scope to spread and transmission of *Salmonella enterica* serovars in the agricultural food chain in the wet markets as most products are sold at ambient temperature and exposed to the environment (Sripaurya et al., 2019). A preceding study stated the incidence of *Salmonella* in various sites of the wet markets indicated a cause of cross-contamination in meat during selling through food contact surfaces or equipment (Nidaullah et al., 2017). Based on the importance of foodborne *Salmonella* at wet markets, this study was aimed to determine the pathogenicity and antimicrobial susceptibility profile of *Salmonella enterica* serovars isolated from poultry processing environments at wet markets in Dhaka, Bangladesh.

## Materials and Methods

### Study design and sample collection

The study was conducted in the 29 chicken wet markets around Dhaka city, the capital of Bangladesh from February to December 2019 in a cross-section manner (Fig 1). Dhaka city is called the biggest chicken selling hub due to the mass population density and economic sovereignty of the population. The sample size was calculated by using the “sample size calculator for prevalence studies, version 1.0.01” based on 25% prevalence of *Salmonella* spp. reported previously in Bangladesh (Naing et al., 2006; Daniel, 1999). The desired individual sample number should not less than 289. Three types of poultry processing environmental samples consisting of carcass dressing water (CDW), chopping board swabs (CBS) and knife swabs (KS) were collected independently as 290. The sterile cotton swabs contained in 10 ml buffered peptone water (BPW) were used for swabbing the samples. The samples were collected aseptically and immediately brought to the Antimicrobial Resistance Action Centre (ARAC) with an insulated icebox. This study received ethical approval from the Ethical Committee of the Animal Health Research Division at the Bangladesh Livestock Research Institute (BLRI), Dhaka, Bangladesh (ARAC: 15/10/2019:05).

**Fig 1.**
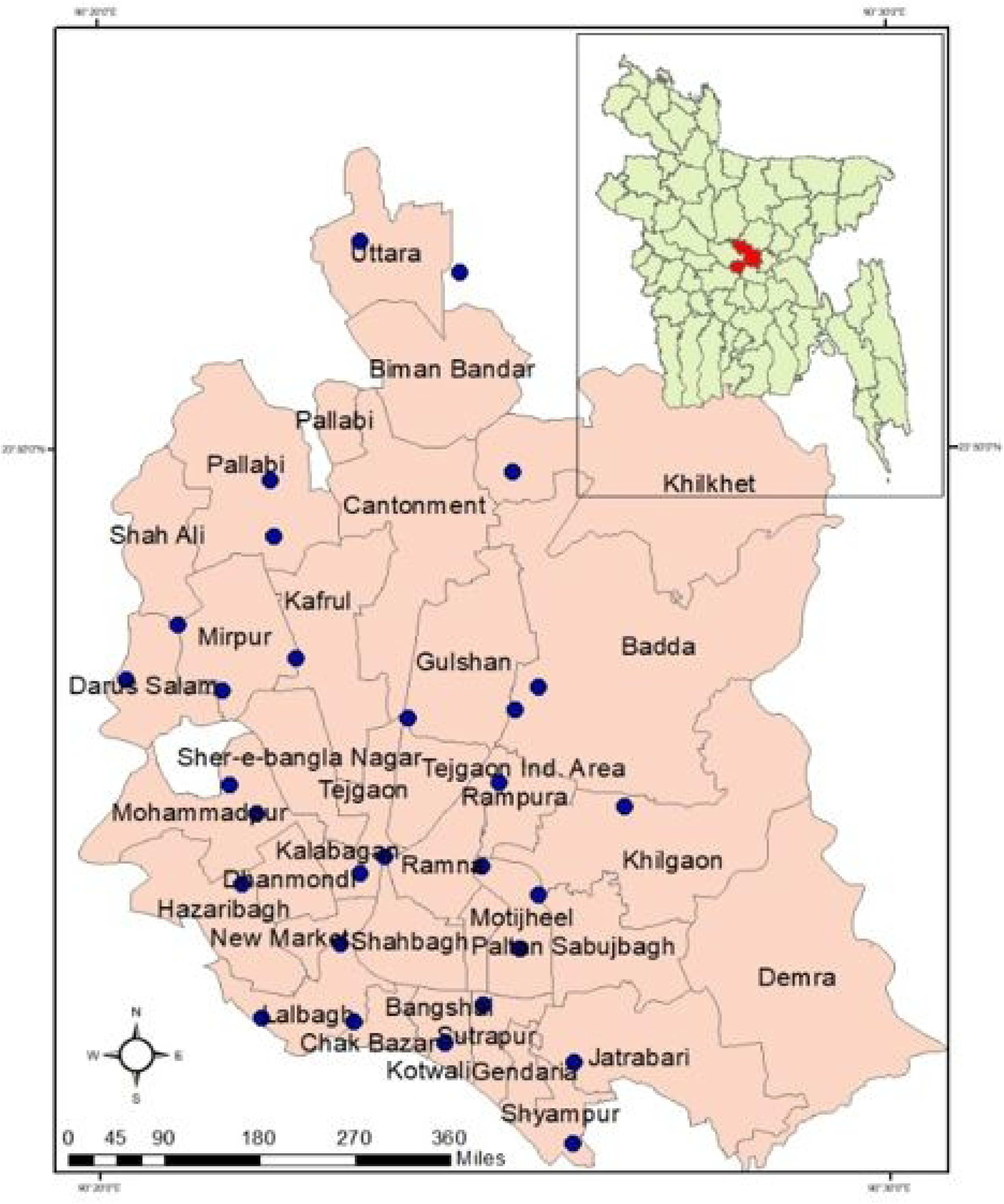
Map of Dhaka city with location of 29 wet markets

### *Salmonella* isolation and identification

*Salmonella* isolation and identification were carried out according to the guidelines of ISO (ISO, 6579:2002) as follows; pre-enrichment of the swab smear in BPW (Oxoid, UK) followed by aerobic incubation at 37°C for 18-24 h. Further, 0.1 mL of the pre-enriched sample was positioned discretely into three different locations on Modified Semisolid Rappaport Vassiliadis (MSRV; Oxoid, UK) agar and incubated at 41.5°C for 20-24 h. Then, a single loop from MSRV was streaked onto Xylose Lysine Deoxycholate (XLD; Oxoid, UK) and another loop onto MacConkey agar (Oxoid, UK) plates and incubated at 37°C for overnight. Typical black centered with reddish zone colony on XLD and colorless colony on MacConkey were picked up and afterward sub cultured on nutrient agar (NA; Oxoid, UK) and screened by biochemical tests: triple sugar iron (TSI), motility indole urea (MIU), catalase and oxidase. Final confirmation was done by the Vitek-2 compact analyzer (bioMérieux, France) followed by PCR (Siddiky et al., 2021).

### DNA extraction

The DNA was extracted using conventional boiling method followed by a proven procedure (Sarker et al., 2019; Heidary et al., 2014). Briefly, the isolate was cultured on NA and incubated overnight at 37°C. Few fresh colonies were harvested from overnight culture and suspended in nuclease-free water. Then bacterial suspension was boiled at 99°C for 15 min followed by chilled on ice. Finally, the debris was separated by centrifugation and supernatant was taken as the DNA template for PCR.

### PCR detection of Salmonella and Salmonella enterica serovars

Uniplex PCR (U-1) was performed to detect *Salmonella* species targeting virulence gene *inv*A (Zahraei-Salehi et al., 2006). Multiplex PCR (M-I) was done to detect *S*. Typhimurium and *S*. Enteritidis (Agron et al., 2001; Alvarez et al., 2004). PCR reaction was adjusted in 25 µL mixture containing 2 µL of DNA template, 12.5 µL of 2x master mix (Go Taq Green Master Mix, Promega), 0.5 μL each of forward and reverse primers (10 pmol/μL) and 9.5 µL nuclease-free water. The PCR products were run at 100 V with 500 mA for 30 min in 1.5% agarose gel containing ethidium bromide. A 100bp DNA ladder (Thermo Scientific, USA) was used as a size marker. The primers used to detect *Salmonella* and *Salmonella* e*nterica* serovars are presented in **Table 1**. *S*. Typhimurium ATCC 14028 and *S*. Enteritidis ATCC 13076 were used as a positive control. Consequently, PCR positive *Salmonella* serovars Typhimurium and Enteritidis was further reconfirmed by the Vitek-2 compact analyzer (bioMérieux, France).

**Table 1.**
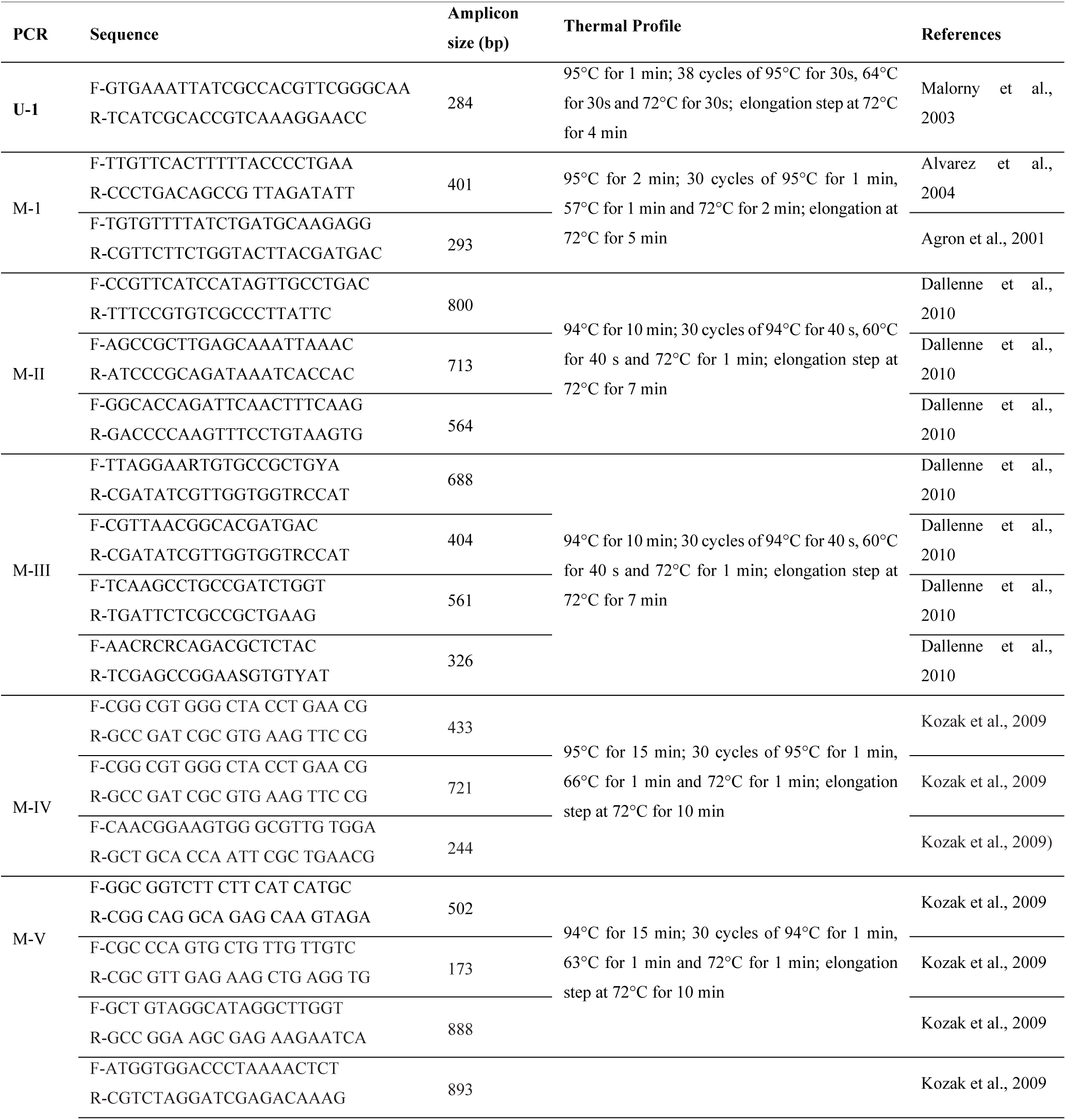
Primers used to detect Salmonella enterica serovars and resistance genes

### Antimicrobial Susceptibility Testing (AST)

AMR profile of all isolates was determined using the Kirby-Bauer disc diffusion method following the guidelines of the Clinical and Laboratory Standards Institute (CLSI, 2019). A panel of 16 antimicrobials representing 10 different classes were selected for AST consisting of aminoglycosides: amikacin (AK, 30µg), gentamicin (CN, 10µg), streptomycin (S, 10µg); carbapenem: meropenem (MEM, 10µg); cephalosporin/beta-lactam antibiotics: ceftriaxone (CRO, 30µg), cefotaxime (CT, 10µg), ceftazidime (CAZ, 30µg), aztreonam (ATM, 30µg); beta-lactamase inhibitors: amoxicillin–clavulanate (AMC, 30µg); penicillins: ampicillin (AMP, 10µg); macrolides: azithromycin (AZM, 15µg); quinolones/fluoroquinolones: ciprofloxacin (CIP, 5µg), nalidixic acid (NA, 30µg); folate pathway inhibitors: sulfamethoxazole-trimethoprim (SXT, 25µg); tetracycline: tetracycline (TE, 10µg); phenicols: chloramphenicol (C, 30µg). The isolates which were resistant to three or more classes of antibiotics were regarded as MDR (Magiorakos et al., 2012). The intermediate isolates were considered as resistant as the acquisition and transition from susceptible to resistance had already begun (Jaja et al., 2019). The positive control was used as *Escherichia coli* ATCC 25922. The multiple antibiotic resistance (MAR) index was calculated and interpreted using the proven method (Adzitey et al., 2012; Titilawo et al., 2015).

### PCR detection of AMR genes

Phenotypically resistant *Salmonella* isolates were screened by PCR for the detection of 14 antibiotic resistance genes comprising of 7 β-lactamase genes (*bla*TEM*, bla*SHV*, bla*OXA*, bla*CTX-M-1*, bla*CTX-M-2*, bla*CTX-M-9 and *bla*CTX-Mg8/25), 3 tetracycline resistant genes (*tet*A*, tet*B and *tet*C), 3 sulfonamide resistant genes (*sul*1*, sul*2 and *sul*3) and single streptomycin resistant gene (*str*A/B). For β-lactam gene, two cycles of multiplex PCR (M-II & M-III) were carried out following the proven method of Dallenne et al. (2010). Consecutively, two cycles of multiplex PCR (M-IV & M-V) were performed to detect the resistance genes for sulfonamide, tetracycline and streptomycin in consistent with established method (Kozak et al., 2009). PCR reaction mixture, as well as gel electrophoresis was done in alignment with the procedures applied for the detection of *Salmonella enterica* serovars in this study. The primers used to detect resistance genes are presented in **Table 1**.

### PCR detection of virulence genes in *Salmonella* isolates

All *Salmonella* isolates were screened for the determination of eight important virulent genes encoding *inv*A, *agf*A, *Ipf*A, *hil*A, *siv*H, *sef*A, *sop*E and *spv*C. The PCR was executed in single reactions following previously used specific primers and thermal profiles (Zahraei-Salehi et al., 2006; Cesco et al 2008; Bäumler &Heffron 1995; Guo et al., 2000; Kingsley et al., 2003; Oliveira et al., 2002; Prager et al., 2003; Swamy et al., 1996). PCR reaction mixture, as well as gel electrophoresis was done in alignment with the procedures applied for the detection of *Salmonella enterica* serovars in this study. The reference positive control (*S*. Typhimurium ATCC 14028 and *S*. Enteritidis ATCC 13076) and negative control (*E. coli* ATCC 25922) were used for validation. The primers used in this study are presented in **Table 2**.

**Table 2.**
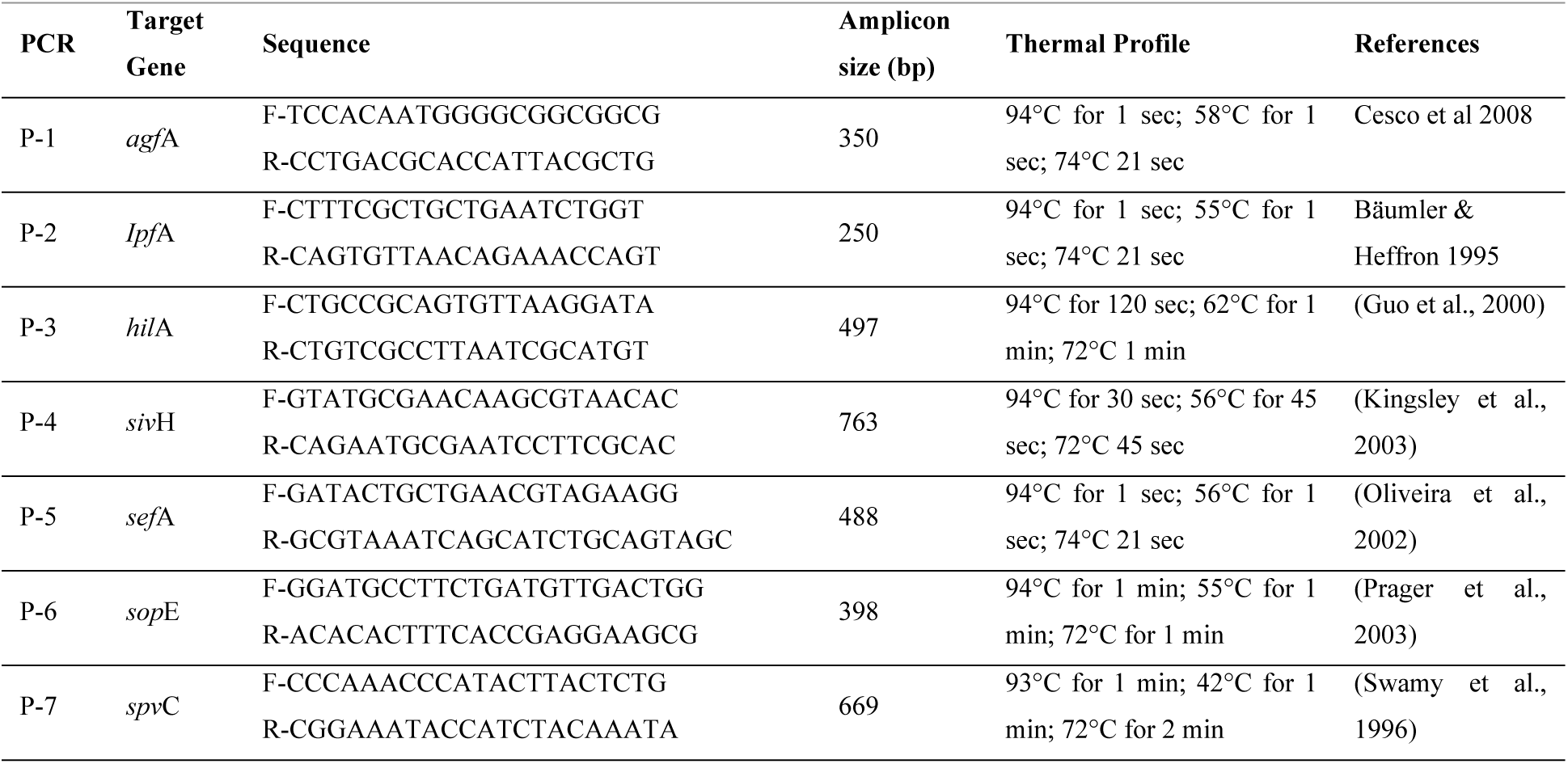
Primers used to detect virulence gene in *Salmonella* isolates

### Statistical analysis

The antimicrobial susceptibility data were presented in Excel sheets (MS-2016) and analyzed by SPSS software (SPSS-24.0). The prevalence was calculated using descriptive analysis and the Chi-square test was applied to determine the level of significance. The statistical significance was determined by *p*-values less than 0.05 (*p* < 0.05).

## Results

### Prevalence of NTS *enterica* serovars

Of all 870 samples, 165 (18.96%) were found to be positive for *Salmonella*. The prevalence of *Salmonella* was found 20% (58 in 290) in CDW, 19.31% (56 in 290) in CBS, and 17.58% (51 in 290) in KS (*p* < 0.05). Meanwhile, the MDR *Salmonella* was found to be 72.41% (42 in 58), 73.21% (41 in 56) and 68.62% (35 in 51) in CDW, CBS, and KS, respectively (*p* < 0.05). The overall prevalence of *S.* Typhimurium, *S.* Enteritidis, and untyped *Salmonella* was found to be 8.96%, 1.6%, and 8.38%, respectively along with an overall MDR were 71.41%. The prevalence of NTS Typhimurium, Enteritidis and untyped *Salmonella* were found to be 7.93%, (23 in 290) 1.72% (5 in 290) and 10.34% (30/290) in CDW, respectively. Likewise, the prevalence of NTS Typhimurium, Enteritidis, and untyped *Salmonella* was found to be 10.34% (30 in 290), 2.06% (6 in 290) and 6.89% (20/290) in CBS, respectively. Consecutively, the prevalence of NTS Typhimurium, Enteritidis, and untyped *Salmonella* were represented 8.62% (25 in 290), 1.03% (3 in 290) and 7.93% (23 in 290) in KS, correspondingly. The prevalence of NTS Typhimurium in CBS were significantly higher compared to CDW and KS (*p* < 0.05).

### Phenotypic resistance patterns of NTS isolates

AST in CDW revealed the highest resistance to ciprofloxacin (68.95%) followed by nalidixic acid (62.06%), tetracycline (60.33%), ampicillin, (58.61%) and streptomycin (56.88%); moderate resistance to gentamicin (39.64%), amoxicillin-clavulanate (31.92%), sulfamethoxazole-trimethoprim, (27.58%) and chloramphenicol (20.67%). On the contrary, low resistance was observed to azithromycin, amikacin and meropenem, respectively. Third-generation cephalosporins (ceftriaxone, cefotaxime, ceftazidime, and aztreonam) were found almost sensitive to all *Salmonella* isolates recovered from CDW (**Fig 2A**). Consecutively, AST of CBS showed higher resistance to streptomycin (64.28%) followed by ciprofloxacin (62.49%), ampicillin (62.27%), tetracycline (60.7%), nalidixic acid (53.56%), and gentamicin (53.56%); moderate resistance (14.27%-46.41%) were recorded to sulfamethoxazole-trimethoprim, amoxicillin-clavulanate, chloramphenicol and azithromycin. Besides, complete sensitive were found in all third-generation cephalosporins including carbapenem (ceftriaxone, cefotaxime, ceftazidime, aztreonam, and meropenem) (**Fig 2B**). Successively, AST of KS exhibited higher resistance to ciprofloxacin (64.69%), ampicillin (64.69%), streptomycin (64.7%), nalidixic acid (58.81%), and tetracycline (54.89%); moderate resistance was recorded to gentamicin (47.05%), sulfamethoxazole-trimethoprim (47.05%), amoxicillin-clavulanate (27.44%) and chloramphenicol (21.56%). On the contrary, very low resistance or almost sensitive were observed to azithromycin, amikacin, meropenem, and third-generation cephalosporins (ceftriaxone, cefotaxime, ceftazidime, and aztreonam) (**Fig 2C**). There was harmony and synergy among the phenotypic resistance patterns of CDW, CBS, and KS. The AST pattern of ciprofloxacin in CDW was significantly higher compared to CBS (*p* < 0.05). Similarly, the AST pattern of gentamicin in CBS were significantly higher compared to CDW and KS (p < 0.05). The highest MAR index value of 0.68, 0.62, and 0.56 was recorded in KS, CDW and CBS, respectively (p < 0.05). The complete sensitive isolates were identified at 1.39% (4 in 290), 2.06% (6 in 290) and 1.03% (3 in 290) in the CDW, CBS, and KS, respectively. The most common antibiotic resistance profile CIP-S-AMP-TE-NA-CN was observed in the *Salmonella* isolates. The detailed phenotypic susceptibility pattern, including *Salmonella enterica* serovars and untyped *Salmonella* from three different sources, is presented in graphical form (**Figs 2A-C**).

**Fig 2.**
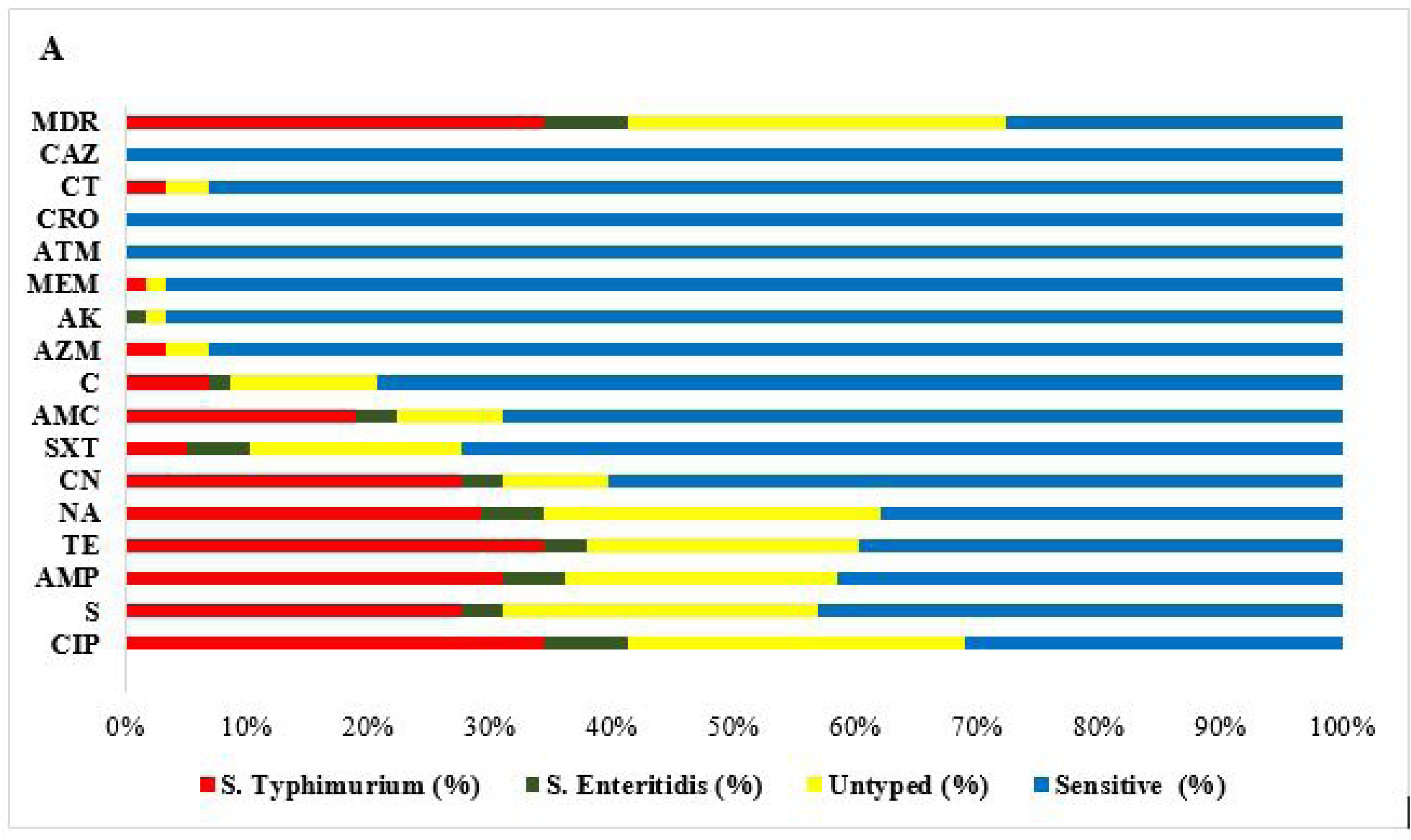
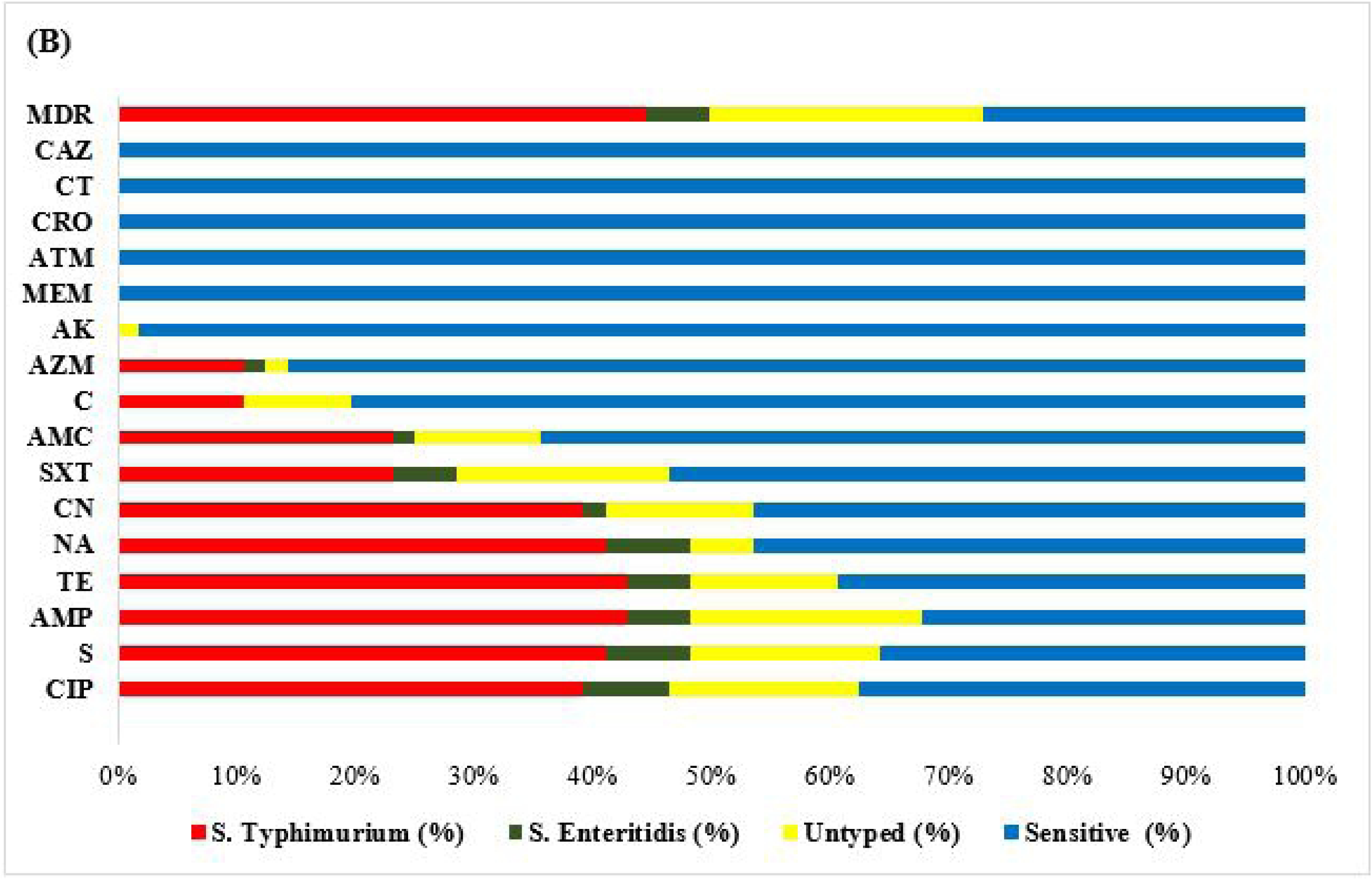
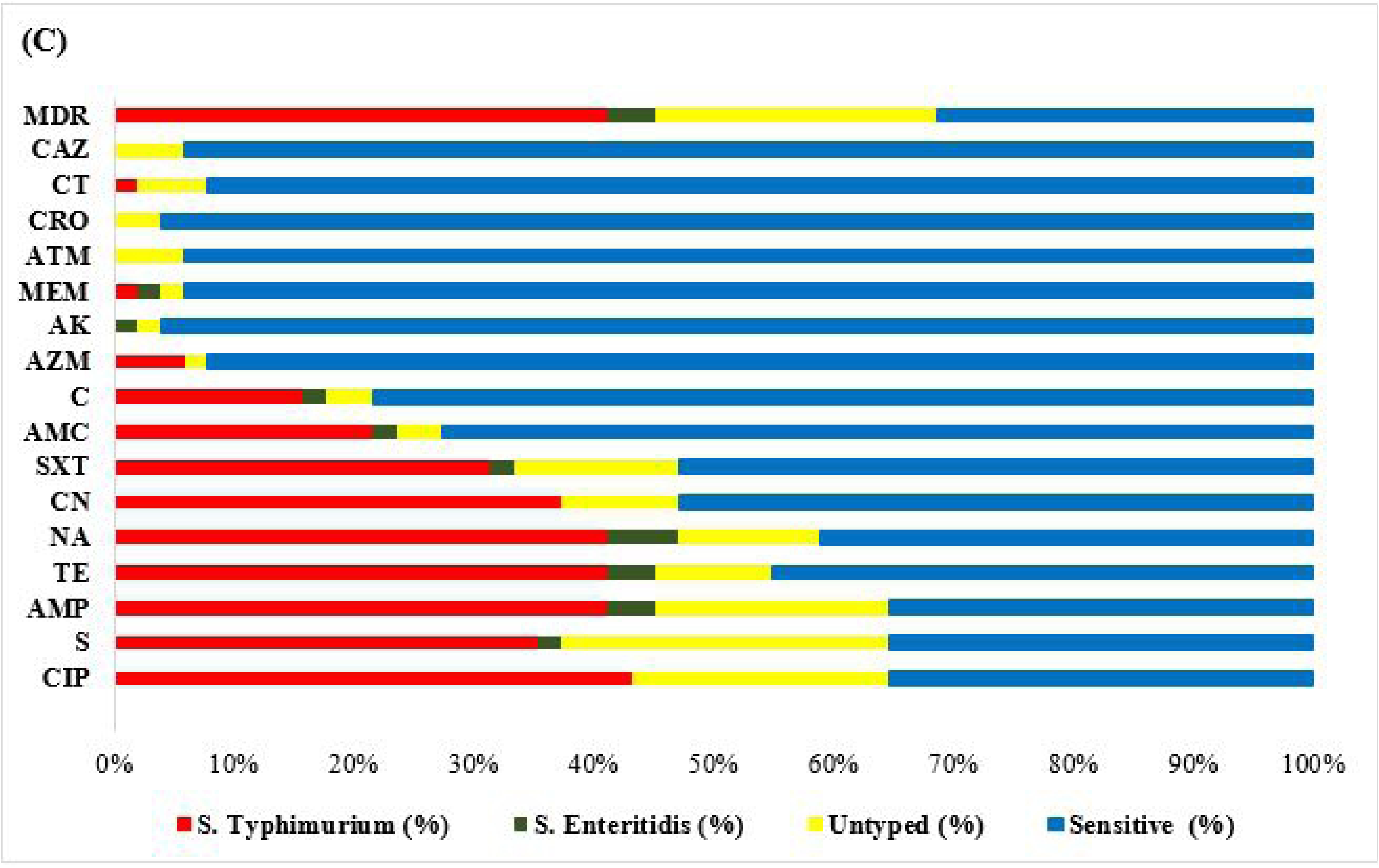
AMR patterns of carcass dressing water (A), chopping board swabs (B) and knife swabs (C).

### Genotypic resistance patterns of NTS isolates

All phenotypically resistant *Salmonella* isolates were screened by PCR for the detection of 14 antibiotic resistant genes encompassing of 7 β-lactamase genes (*bla*TEM*, bla*SHV*, bla*OXA*, bla*CTX-M-1*, bla*CTX-M-2*, bla*CTX-M-9 and *bla*CTX-Mg8/25), 3 tetracycline resistant genes (*tet*A*, tet*B and *tet*C), 3 sulfonamide resistant genes (*sul*1*, sul*2, and *sul*3) and single streptomycin resistant gene (*str*A/B) recovered from CDW (**Fig 3A**), CBS (**Fig 3B**) and KS (**Fig 3C**).

**Fig 3.**
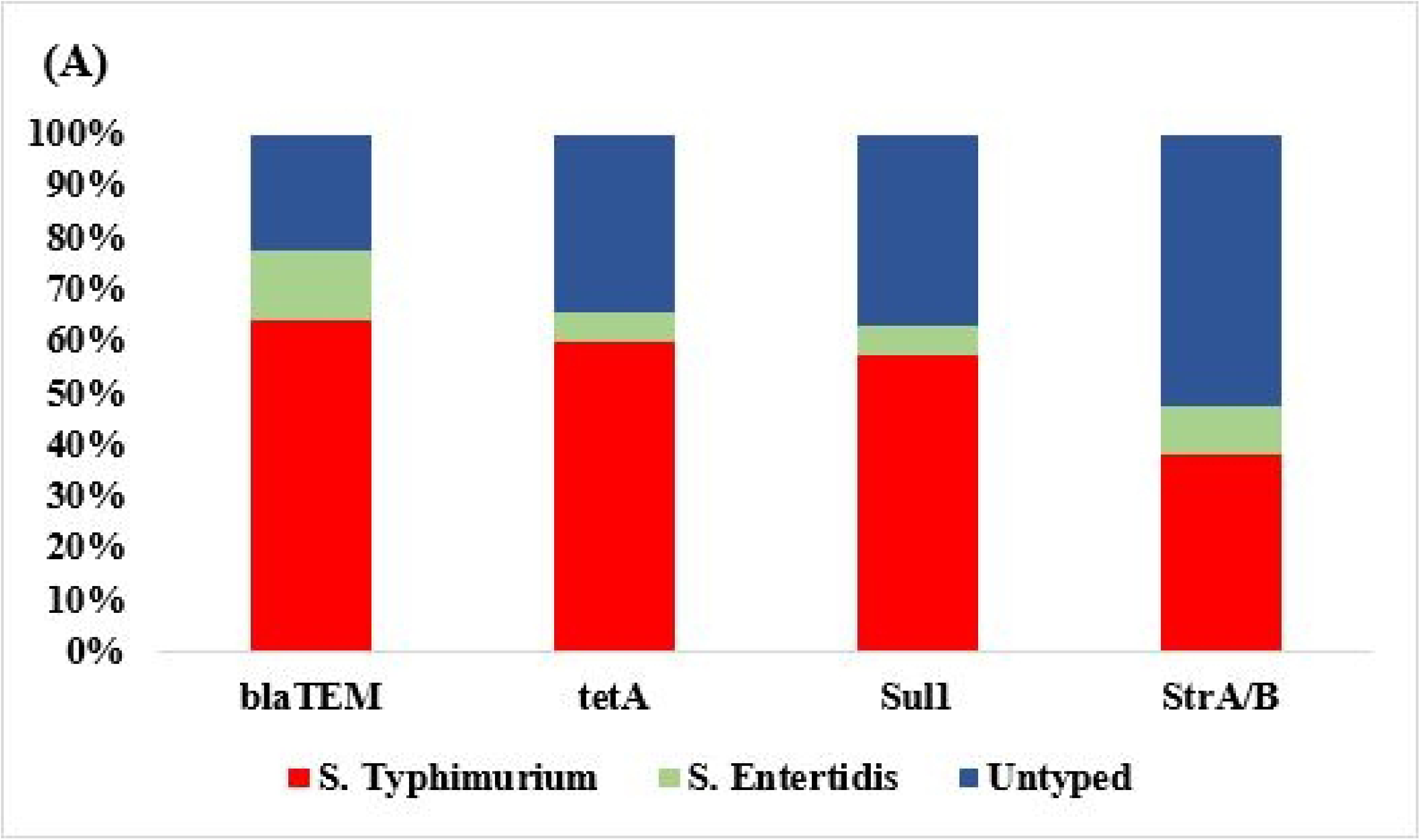
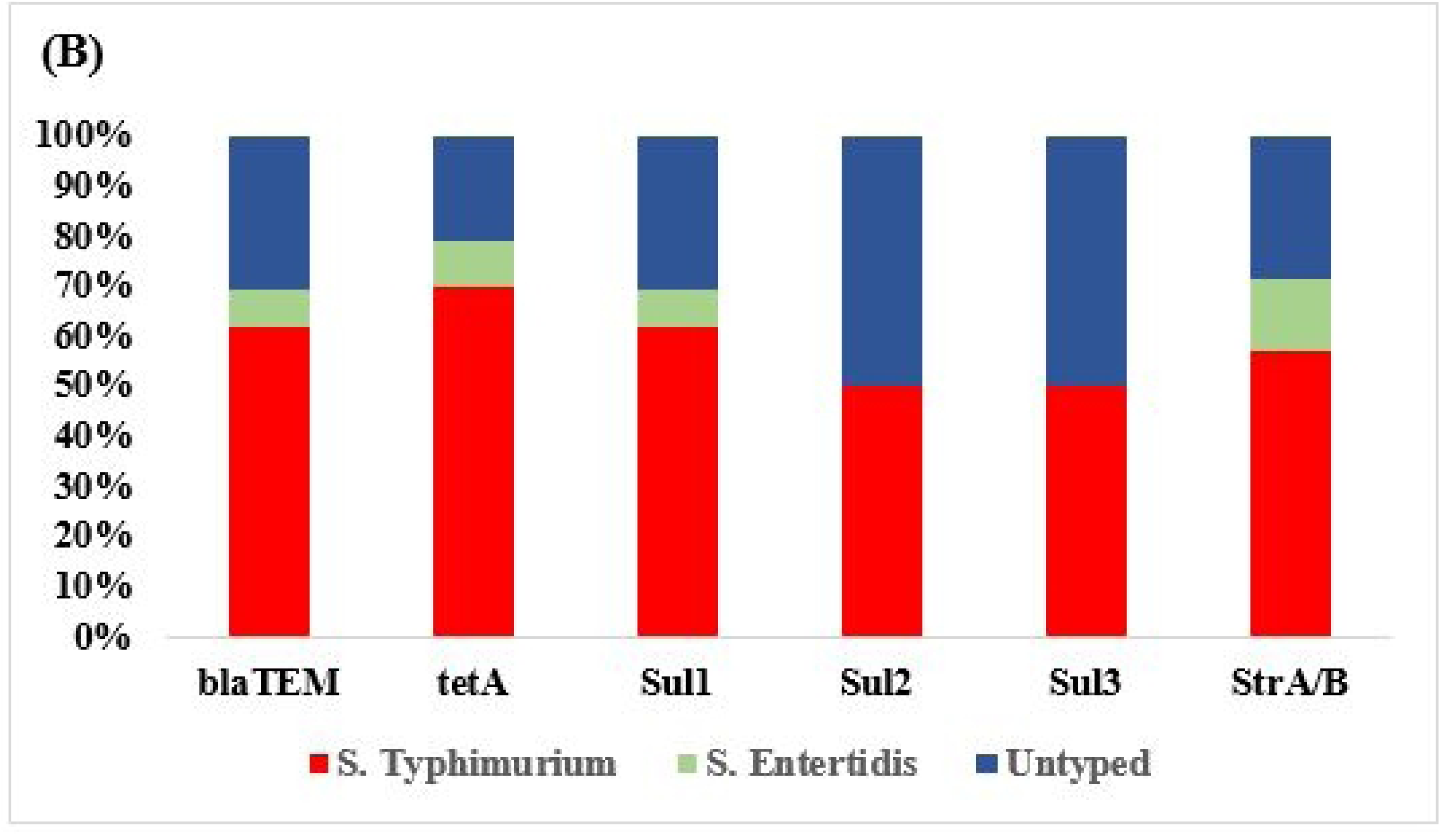
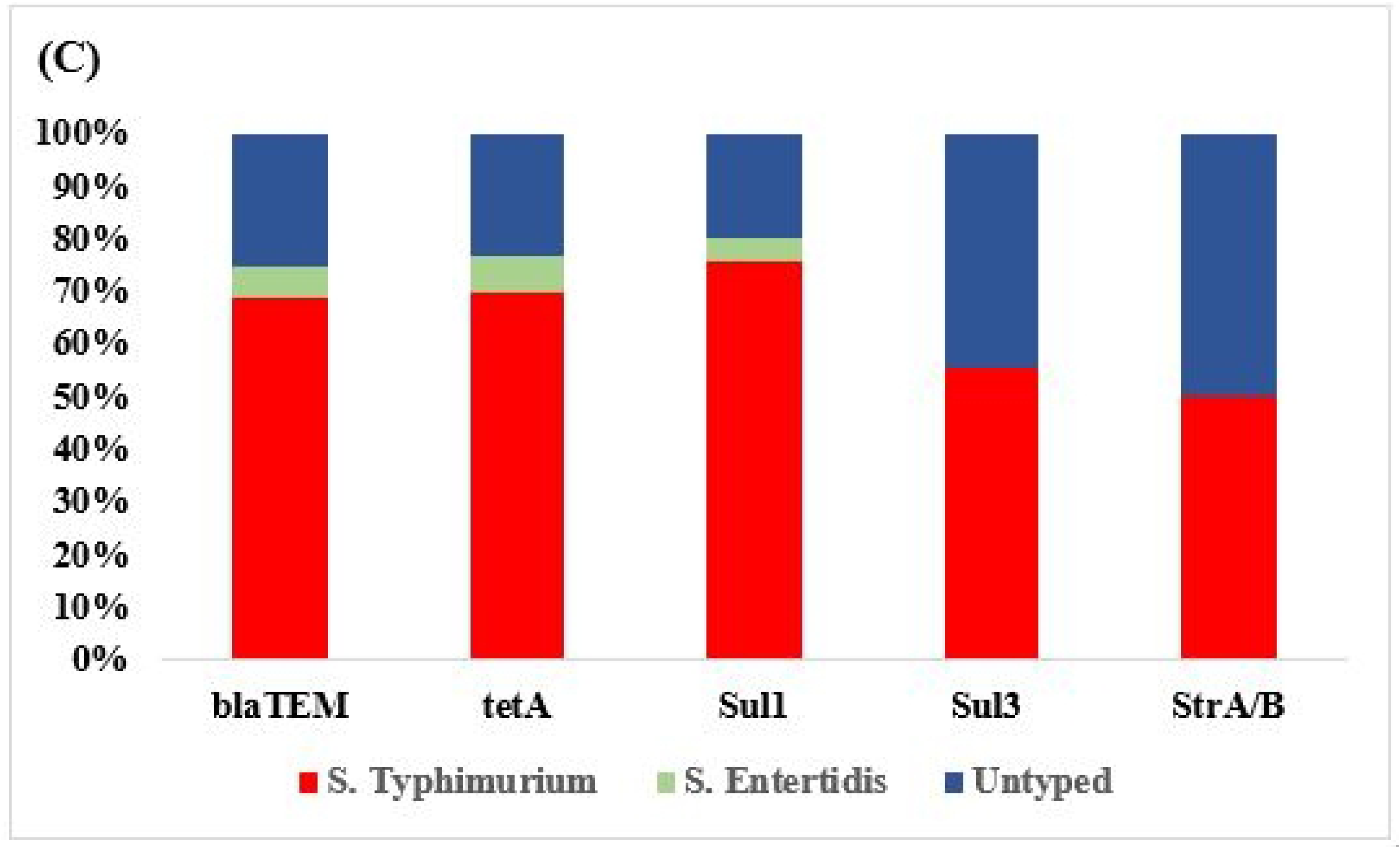
Genotypic resistance patterns of carcass dressing water (A), chopping board swabs (B) and knife swabs (C).

Out of seven, only one ESBL gene, *bla*TEM was detected with a prevalence rate of 62.06%, 69.62% and 62.73% in CDW, CBS, and KS, respectively. Consecutively, out of three tetracycline resistant genes, only one *tet*A was identified with the prevalence level of 60.32%, 58.92%, and 58.81% in CDW, CBS, and KS, respectively. Sequentially, the prevalence of *sul*1 gene were found 60.33%, 69.62%, and 49.01% in CDW, CBS, and KS, respectively. Furthermore, *sul*2 and *sul*3 were also found only in CBS with lower prevalence level of 3.56% and 3.56%, respectively. Similarly, *sul*3 gene was detected only in KS with a prevalence rate of 17.62%. Moreover, the streptomycin resistance gene, *str*A/B was detected with a prevalence rate of 36.2%, 24.99%, and 31.36% in CDW, CBS and KS, respectively. The detailed genotypic susceptibility pattern, including *Salmonella enterica* serovars and untyped *Salmonella* from three different sources, is presented in graphical form (**Figs 3A-C**). The *sul*1gene in CBS were significantly higher compared to CDW and KS (*p* < 0.05). Similarly, *str*A/B gene in CDW was significantly higher compared to KS and CBS (*p* < 0.05). There was a strong correlation and synergy persists among the genotypic features.

### PCR detection of virulence genes for NTS isolates

All *Salmonella isolates* were screened by PCR to monitor eight common virulence genes namely *inv*A, *agf*A, *Ipf*A, *hil*A, *siv*H, *sef*A, *sop*E, and *spv*C. *S*. Enteritidis and untyped *Salmonella* isolates were found positive for all eight common virulence genes whereas *S.* Typhimurium harbored six virulence genes (**Fig 4**). There was a strong correlation exist among the virulence genes in CDW, CBS, and KS (*p* < 0.05).

**Fig 4.**
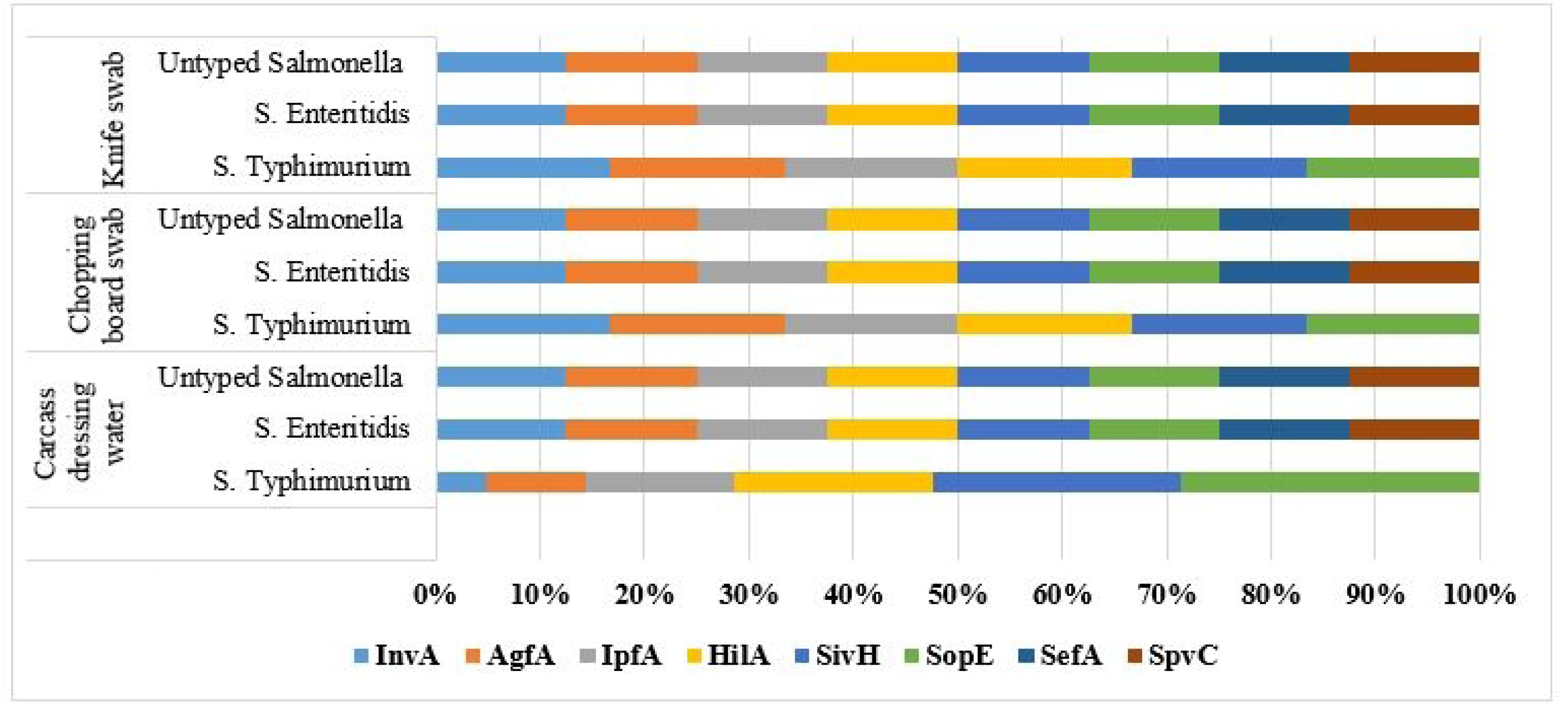
Virulence genes distribution among *Salmonella* isolates

## Discussion

Very few studies have reported NTS *enterica* serovars isolated from poultry processing environments at wet markets in Bangladesh. In our study, we found the overall prevalence of *Salmonella* 18.96% in poultry processing environmental samples including CDW, CBS and KS. Previously, the prevalence of *Salmonella* in Bangladesh was found 23.33% in poultry slaughter specimens (Momtaz et al., 2018); 26.6% in chicken cloacal swab, intestinal fluid, egg surface, hand wash, and soil of chicken market samples (Akond et al., 2012); 25.35% in the chicken cloacal swab, eggshells, intestinal contents, liver swabs, broiler meat, and swabs of slaughterhouse (Karim et al., 2017); 35% in broiler farms settings (Alam et al., 2020); 23.53% in poultry samples (Al Mamun et al., 2017); 37.9% in poultry production settings (Mahmud et al., 2011); 31.25% in broiler farm settings (Mridha et al., 2020); 42% in broiler chicken (Sarker et al., 2021) and 65% in frozen chicken meat (Parvin et al., 2020). Successively, Siddiky et al. (2021) found the prevalence of *Salmonella* 8.62% in broiler, 6.89% in sonali and 3.1% in native chicken cecal contents. The prevalence of *Salmonella* isolates in our findings was compatible with previous data in Bangladesh.

Continually, the overall prevalence of *Salmonella* was found at 17.3% in butcher shops in Ethiopia. They identified *Salmonella* in KS, CBS, hand washings, and meat which has strong synergy with our findings (Garedew et al., 2015). Rather, the wet market based study in India revealed the prevalence of *Salmonella* 14.83% in chicken meat shops (Sharma et al., 2019); 19.04% in retail chicken shops (Waghamare et al., 2017), and 23.7% in white and red meat at local markets (Kaushik et al., 2014). A study demonstrated the high prevalence of *Salmonella* (88.46%) in poultry processing and environmental samples obtained from wet markets and small-scale processing plants in Malaysia (Nidaullah et al., 2017). Rusul et al. (1996) reported the prevalence of *Salmonella* 35.5% and 50.0% in broiler carcasses at the wet markets and processing plants, respectively. Furthermore, the overall prevalence of *Salmonella* serovars identified 23.5% in ducks, duck rearing, and processing environments in Penang, Malaysia (Adzitey et al., 2012). Moreover, the recent prevalence of *Salmonella* at home and abroad was linked to our findings. In our study, the prevalence of NTS Typhimurium was found to be 7.93%, 10.34%, and 8.62% respectively in CDW, CBS and KS. Consecutively, the occurrence of *S.* Enteritidis were found 1.72%, 2.06%, and 1.03% in CDW, CBS, and KS respectively. Siddiky et al. (2021) found the overall prevalence of *S*. Typhimurium and *S*. Enteritidis at the rate of 3.67% and 0.57% in chicken cecal contents. Some correlation of the prevalence of *Salmonella enterica* serovars between caecal content and environmental samples was observed. Successively, Thung et al. (2018) who found *S*. Enteritidis, and *S*. Typhimurium with prevalence rate of 6.70%, and 2.50%, respectively in raw chicken meat at retail markets in Malaysia. In our study, *S*. Typhimurium was the major serovar which was consistent with the findings of McCrea et al. (2006) who identified *S*. Typhimurium as the major *Salmonella* serovar from a poultry market in California, USA. Saitanu et al. (1994) also reported *S*. Typhimurium (5.5%) was the most predominant serotype in duck eggs in Thailand. The study conducted in Bangladesh revealed *S*. Typhimurium incidence 15.91% in the broiler production systems (Islam et al., 2016); 85% in broiler farm samples (Alam et al., 2020) and 5% in commercial layer farm settings (Parvej et al., 2016). The higher occurrence of *S*. Typhimurium in our study is synchronized with findings at home and abroad. Consecutively, *S*. Typhimurium and *S*. Enteritidis were isolated from raw chicken meat at retail markets in Malaysia (Thung et al., 2018); higher prevalence of *S*. Enteritidis (21.9%) and *S*. Typhimurium (9.4%) were isolated from chicken in Turkey (Arkali et al., 2020); *Salmonella enterica* serovars were identified in backyard poultry flocks in India (Samanta et al., 2014); *S*. Enteritidis and *S*. Typhimurium recovered from chicken meat in Egypt (Tarabees et al., 2017). Similarly, Suresh et al. (2006) recovered *S*. Typhimurium and *S*. Enteritidis in high proportion compared to other serovars from various poultry products in India. Furthermore, China and some European countries detected *S*. Enteritidis and *S*. Typhimurium as the most prevalent serotypes (Osimani et al., 2016; Zeng et al., 2019).

In our study, NTS *enterica* serovars Typhimurium and Enteritidis along with untyped *Salmonella* were found higher resistance to ciprofloxacin, streptomycin, gentamicin, ampicillin, tetracycline, and nalidixic acid; moderate resistance to sulfamethoxazole-trimethoprim, amoxicillin-clavulanate, chloramphenicol, and azithromycin. Alam et al. (2020) identified a high level of resistance (77.1% to 97.1%) of *Salmonella* isolates to tetracycline, ampicillin, streptomycin, and chloramphenicol. Further, Parvin et al. (2020) reported the highest resistance of *Salmonella* isolates against oxytetracycline (100%), followed by trimethoprim-sulfamethoxazole (89.2%), tetracycline (86.5%), nalidixic acid (83.8%), amoxicillin (74.3%), and pefloxacin (70.3%). Successively, Mridha et al. (2020) demonstrated higher to moderate resistance of *Salmonella* isolates to erythromycin, tetracycline, amoxicillin, and azithromycin. Sequentially, Sobur et al. (2019) found higher resistance of *Salmonella* isolates to tetracycline, ciprofloxacin, and ampicillin. Continually, it has been observed that *Salmonella* originated from fecal samples of chickens, ducks, geese and pigs were resistant to nalidixic acid (48.8%), tetracycline (46.9%), ampicillin (43.2%), streptomycin (38.3%), and trimethoprim/sulfamethoxazole (33.3%) respectively (Long et al., 2016; Im et al., 2015; Vuthy et al., 2017). Concurrently, high resistance to ciprofloxacin (77%), sulfisoxazole (73%) and ampicillin (55.6%) was recorded in chicken hatcheries in China (Xu et al., 2020); tetracycline and ampicillin were found highly resistance in wet markets, Thailand (Sripaurya et al., 2018); higher resistance to nalidixic acid (99.5%), ampicillin (87.8%), tetracycline (51.9%), ciprofloxacin (48.7%), and trimethoprim-sulfamethoxazole (48.1%) were found in broiler chickens along the slaughtering process in China (Zhu et al., 2017); *S*. enterica serovar Typhimurium was found resistant to ampicillin, tetracycline, and sulphamethoxazole isolated from chicken farms in Egypt (El-Sharkawy et al., 2017); high resistance to ampicillin (95.71%), ciprofloxacin (82.86%), tetracycline (100%), and nalidixic acid (98.57%) was observed in isolated NTS at retail chicken meat stores in northern India (Sharma et al., 2019). Furthermore, Siddiky et al. (2021) reported the highest resistance pattern of *S*. Typhimurium isolates to ciprofloxacin (100%) and streptomycin (100%) followed by tetracycline (86.66%), nalidixic acid (86.66%), gentamicin (86.66%), ampicillin (66.66%) and amoxicillin–clavulanate (40%) in broiler chicken. Sequentially, Siddiky et al. (2021) identified maximum resistance of *S*. Enteritidis isolates to streptomycin (100%) followed by ciprofloxacin (80%), tetracycline (80%), gentamicin (80%), and moderate resistance to amikacin (20%), amoxicillin–clavulanate (20%), azithromycin ((20%), and sulphamethazaxole-trimethoprim (20%) in broiler chicken. There was a very strong correlation and alignment was observed at home and abroad with phenotypic resistance patterns.

In our study, out of seven, a single beta-lactam-resistant *bla*TEM gene was detected in all three types of samples with a diverse prevalence rate. The prevalence of *bla*TEM was detected at 62.06%, 69.62%, and 62.73 in CDW, CBS, and KS. Ahmed et al. (2014) detected a higher prevalence of *bla*TEM gene mediated ESBL production among *Salmonella* isolated from humans in Bangladesh. Yang et al. (2010) identified *bla*TEM, a gene encoded for beta-lactamases resistance, in 51.6% resistant *Salmonella* isolates. Aslam et al. (2012) reported *bla*TEM gene in *Salmonella* isolated from retail meats in Canada with a 17% prevalence rate. Lu et al. (2011) detected only 81.2% *bla*TEM gene, while *bla*CTX-M could not be detected in any of the examined isolates. Similarly, Van et al. (2008) found only the *bla*TEM gene in *E.* coli recovered from raw meat and shellfish in Vietnam. The emergence of *bla*TEM mediated ESBL producing NTS *enterica* serovars indicated the use of beta-lactam antibiotics in poultry farming practices. Siddiky et al. (2021) detected only the *bla*TEM gene from chicken cecal contents and Xiang et al. (2020) reported plasmid-borne and easily transferable *bla*OXA-1 and *bla*TEM-1 genes. Consecutively, Suresh et al. (2019) detected *bla*TEM as the predominant gene in food of animal origin in India. The results of the *bla*TEM gene were in agreement and relevant to previous observations. Successively, in our study, out of three only a single tetracycline resistance gene *tet*A was found with the prevalence rate of 60.32%, 58.92%, and 58.81% in carcass dressing water, chopping board swab, and knife swab respectively. In addition, *sul*1, *sul*2, and *sul*3 genes were detected in NTS *enterica* serovars, while *sul*1 was predominant. Likewise, streptomycin resistance gene (*str*A/B) was isolated in serovars of NTS with reliable prevalence rate. Arkali and Çetinkaya, (2020) detected 58% positive *sul*1 gene from the *Salmonella* isolates of chickens in eastern Turkey. Consequently, Jahantigh et al. (2020) detected the most prevalent *tet*A gene from broiler chicken in Iran. Successively, Vuthy et al. (2017) discovered *bla*TEM, *tet*A and *str*A/B gene from chicken food chain while Sin et al. (2020) isolated *tet*A and *sul*1 gene from chicken meat in Korea. Zhu et al. (2017) who isolated beta lactam (*bla*TEM), tetracycline resistant (*tet*A, *tet*B, *tet*C) and sulfonamide resistant (*sul*1, *sul*2 and *sul*3) gene with prevalence at 94.6%, 85.7%, and 97.8% in *Salmonella* isolated from slaughtering process in China. El-Sharkawy et al. (2017) who revealed *bla*TEM, *tet*A, *tet*C, *sul*1, and *sul*3 gene from *S*. Enteritidis isolates at a chicken farm in Egypt. Doosti et al. (2016) detected *str*A/B (37.6%) from *S*. Typhimurium isolates at poultry carcasses in Iran. Sharma et al. (2019) detected the most predominant *tet*A and *bla*TEM gene in NTS isolated from retail chicken shops in India. Continually, Alam et al. (2020) revealed *tet*A (97.14%) and *bla*TEM-1 (82.85%) gene in broiler farms whilst Siddiky et al. (2021) detected *tet*A, *sul*1, and *str*A/B gene from chicken cecal contents in Bangladesh. The genotypic resistance patterns were well matched with former findings at home and abroad.

In our study, harmonic and proportioned correlation were existent between genotypic and phenotypic resistance decoration. These findings were in agreement and alignment with the observations of previous study conducted across the globe (Alam et al., 2020; Zhu et al., 2017; Siddiky et al., 2021). However, some disagreement between phenotypic and genotypic resistance pattern might be due to misalignment of antibiotic disk, sensitivity and specificity of disk, primers, concentration of inoculum, laboratory capacity and individual skills (Siddiky et al., 2021). Rather, some research findings supported the misalignment between genotypic and phenotypic resistance patterns (Van et al., 2008; Paterson et al., 2005).

In our study, MDR *Salmonella* embedded mostly with ciprofloxacin, streptomycin, tetracycline, ampicillin, gentamicin and nalidixic acid probably due to common and frequent uses of these antibiotics in poultry production settings in Bangladesh (Al Masud et al., 2020; Parvej et al., 2016). CLSI (2019) reported *Salmonella* became natural resistant to first and second-generation cephalosporin and aminoglycosides. MDR along with higher resistance to ciprofloxacin is very much alarming in human treatment as WHO recommended ciprofloxacin, a first-line drug treatment of intestinal infections. Besides, watch group antibiotic ciprofloxacin had higher resistance and azithromycin had moderate resistance reflects the severity of the resistance pattern of *Salmonella* serovars (WHO, 2019). Furthermore, more resistance to tetracycline indicated the massive use of therapeutic and growth enhancers in poultry production (Wang et al., 2013). The emergence of *bla*TEM mediated ESBL producing *Salmonella enterica* serovars indicated the use of beta-lactam antibiotics in the poultry production cycle. Moreover, ESBL is usually encoded by large plasmids that are transferable from strain to strain and between bacterial species (Ahmed et al., 2014; González et al., 2013).

Virulence gene analysis indicated *S*. Enteritidis and untyped *Salmonella* isolate carried eight virulence genes including two types of *Salmonella* pathogenicity islands (SPI-1 and SPI-2) and many adhesion-related virulence genes. Virulence genes along with MDR resistance pattern would accelerate the infectivity of *Salmonella* isolates (Huehn et al., 2010). The emergence of antibiotic resistance of *Salmonella* isolates depends on its genetic and pathogenicity mechanisms which can enhance the survivability by preserving drug resistance genes (Aslam et al., 2012). The virulence gene was found more prevalent in *S.* Enteritidis and untyped *Salmonell*a isolates compared to *S*. Typhimurium isolates. Six common virulence genes (*inv*A, *agf*A, *Ipf*A, *hil*A, *siv*H, and <u>spv</u>C) were detected in all isolates of *Salmonella* which was incompatible with prior findings around the world (Amini et al., 2010; Borges et al., 2013; Campioni et al., 2012; Crâciuna et al., 2012). Moreover, *sef*A and *spv*C genes were detected in *S*. Enteritidis and untyped *Salmonella* isolates whilst none *S.* Typhimurium isolates carried *sef*A and *spv*C genes. Alike findings have been recorded formerly by researchers (Borges et al., 2013). The higher occurrence of *sef*A in *S*. Enteritidis was compatible with prior findings (Amini et al., 2010; Crâciuna et al., 2012), and *sef*A was somewhat considered target gene to encounter *S*. Enteritidis through the PCR method (Amini et al., 2010). Successively, the *inv*A was the most common and virulent gene present in all *Salmonella* isolates and considered as a target gene to identify *Salmonella* species (Nayak et al., 2004; Elkenany et al., 2009). Continually, the *hil*A gene played a key role to exaggerate *Salmonella* virulence by stimulating the expression of invasion into the cell (Ramatla et al., 2017; Cardona-Castro et al., 2002). Furthermore, the virulence gene *inv*A and *hil*A could be considered target genes to rapid and reliable detection of *Salmonella* through PCR method. The higher occurrence of *lpf*A, *agf*A, and *sop*E were consistent with previous research results (Collinson et al., 1993; Borsoi et al., 2009). The occurrence of the *sop*E gene (100%) in *S*. Enteritidis was correlated with earlier studies (Batchelor et al., 2005). Further, *agf*A gene liable for biofilm development along with adhesion to cells during infection process (Yoo et al., 2013). In our study, the plasmid-mediated *spv*C virulence gene was detected in *S*. Enteritidis and untyped *Salmonella* isolates which have similarities to earlier observations (Amini et al., 2010; Castilla et al., 2016; Motta et al., 2003). It was previously found *S*. Enteritidis had 92% *spv*C gene while *S*. Typhimurium had only 28% and *S.* Hadar had no (Borges et al., 2013). The higher prevalence of major virulence genes indicated the pathobiology as well as the public health implications of the serovars. Moreover, all *Salmonella enterica* isolates were found to be highly invasive and enterotoxigenic, which had a significant public health impact. This was the first-ever attempt to determine a wider range of *Salmonella* virulence genes from poultry processing environments in wet markets in Dhaka, Bangladesh.

Our results demonstrated that wet markets where chicken has been slaughtered and processed that could spread and harbour NTS *enterica* serovars. Many studies have shown that cross-contamination of poultry could occur during processing and skinning in wet markets due to poor sanitary and hygienic measures (Nidaullah et al., 2017). In wet markets, the sources of contamination may be vendor, chopping board swab, knife swab, carcass dressing water, defeathering machines, scalding water, tanks, floors, drains and work benches etc. (Nidaullah et al., 2017). Free roaming MDR NTS serovars in poultry processing environments could facilitate release into the food chain, agricultural goods, and human populations in wet markets. Furthermore, *Salmonella* serovars were able to survive longer with soil-formed biofilms, and these biofilms were protected from detergents and sanitizers. Therefore, cleaning and sterilization of the knife, chopping board and frequent change of carcass dressing water is crucial to reduce the burden of horizontal transmission of *Salmonella* enterica in wet markets. The root causes indicated that MDR and highly pathogenic NTS *enterica* has been emerged in poultry due to the irrational use of antimicrobials in farming practices.

## Conclusion

The higher prevalence of multiple profiles of virulence and multidrug resistance genes of NTS *enterica* serovars in poultry processing environments drew public health attention. The chicken carcass was dressed and processed in open environments in wet markets exacerbated the spread of the pathogen. The unclean and utilized utensils such as chopping boards and knives were used for chicken processing and cutting. Successively, unclean and dirty water were used frequently for chicken carcass dressing and washing. Their numerous risk factors are prevailing at wet markets that can trigger the spread and contamination of NTS serovars. The wet market can be considered as the hotspots of harboring NTS serovars which can easily anchor in the food chain as well as human health. The hidden sources of these MDR pathogens were undoubtedly from chicken which indicated more therapeutic and preventive exposure of antimicrobials during the production cycle. The clean, hygienic and ambient poultry processing environments with good carcass processing practices might reduce the spread and contamination at wet markets. Besides, prudent and judicious uses of antimicrobials have to be followed in farming practices as poultry farming is considered the fertile ground for uses of antimicrobials. This study could address the potential risk associated with the spread of NTS with multidrug resistance to humans as well as highlight the need of implementing a strict hygiene and sanitation standard in local wet markets.

## Supporting Information

**S1 Text: Phenotypic and genotypic antimicrobial resistance data**

## Acknowledgments

This work was supported by the project on “combating the threats of antimicrobial resistance and zoonotic diseases to achieve the GHSA in Bangladesh”. The author highly acknowledge the logistic, financial and technical support and cooperation of Director, Antimicrobial Resistance Action Centre, Bangladesh Livestock Research Institute.

## Author Contributions

Conceptualization: Nure Alam Siddiky, Md. Shahidur Rahman Khan, Mohammed A. Samad; Data curation: Nure Alam Siddiky; Md Samun Sarker; Formal analysis: Nure Alam Siddiky; Investigation: Nure Alam Siddiky; Methodology: Nure Alam Siddiky, Md Samun Sarker; Software: Nure Alam Siddiky, Md Samun Sarker; Supervision: Md. Shahidur Rahman Khan, Mohammed A. Samad, Md. Tanvir Rahman, Md. Abdul Kafi; Validation: Md. Tanvir Rahman, Md. Abdul Kafi; Visualization: Nure Alam Siddiky, Md Samun Sarker; Writing-original draft: Nure Alam Siddiky, Md Samun Sarker; Writing-review & editing: Md. Shahidur Rahman Khan, Mohammed A. Samad, Md. Tanvir Rahman, Md. Abdul Kafi.

## References

Adzitey F, Rusul G, Huda N. Prevalence and antibiotic resistance of Salmonella serovars in ducks, duck rearing and processing environments in Penang, Malaysia. Food Res Int. 2012 Mar 1; 45(2):947–52. https://doi.org/10.1016/j.foodres.2011.02.051

Agron PG, Walker RL, Kinde H, Sawyer SJ, Hayes DC, Wollard J, Andersen GL. Identification by subtractive hybridization of sequences specific for Salmonella enterica serovar Enteritidis. Appl Environ Microbiol. 2001 Nov 1; 67(11):4984–91. https://doi.org/10.1128/aem.67.11.4984-4991.2001

Ahmed D, Ud-Din AI, Wahid SU, Mazumder R, Nahar K, Hossain A. Emergence of blaTEM type extended-spectrum β-lactamase producing *Salmonella* spp. in the urban area of Bangladesh. Int Sch Res Notices. 2014; 2014. https://doi.org/10.1155/2014/715310

Akond MA, Shirin M, Alam S, Hassan SM, Rahman MM, Hoq M. Frequency of drug resistant *Salmonella* spp. isolated from poultry samples in Bangladesh. Stamford J Microbiol. 2012; 2(1):15–9. https://doi.org/10.3329/sjm.v2i1.15207

Al Mamun MA, Kabir SL, Islam MM, Lubna M, Islam SS, Akhter AT, et al. Molecular identification and characterization of *Salmonella* species isolated from poultry value chains of Gazipur and Tangail districts of Bangladesh. Afr J Microbiol Res. 2017 Mar 21; 11(11):474–81. https://doi.org/10.5897/AJMR2017-8431

Al Masud A, Rousham EK, Islam MA, Alam MU, Rahman M, Al Mamun A, et al. Drivers of antibiotic use in poultry production in Bangladesh: dependencies and dynamics of a patron-client relationship. Front Vet Sci. 2020; 7. https://doi.org/10.3389/fvets.2020.00078

Alam SB, Mahmud M, Akter R, Hasan M, Sobur A, Nazir KH, Noreddin A, Rahman MT, El Zowalaty ME, Rahman M. Molecular detection of multidrug resistant *Salmonella* species isolated from broiler farm in Bangladesh. Pathogens. 2020 Mar; 9(3):201. https://dx.doi.org/10.3390%2Fpathogens9030201

Alvarez J, Sota M, Vivanco AB, Perales I, Cisterna R, Rementeria A, Garaizar J. Development of a multiplex PCR technique for detection and epidemiological typing of *Salmonella* in human clinical samples. J Clin Microbiol. 2004 Apr 1; 42(4):1734–8. https://dx.doi.org/10.1128%2FJCM.42.4.1734-1738.2004

Amini K, Salehi TZ, Nikbakht G, Ranjbar R, Amini J, Ashrafganjooei SB. Molecular detection of invA and spv virulence genes in *Salmonella* enteritidis isolated from human and animals in Iran. Afr J Microbiol Res. 2010 Nov 4; 4(21):2202–10. https://doi.org/10.5897/AJMR.9000508

Arkali A, Çetinkaya B. Molecular identification and antibiotic resistance profiling of *Salmonella* species isolated from chickens in eastern Turkey. BMC Vet Res. 2020 Dec; 16(1):1–8. https://doi.org/10.1186/s12917-020-02425-0

Aslam M, Checkley S, Avery B, Chalmers G, Bohaychuk V, Gensler G, et al. Phenotypic and genetic characterization of antimicrobial resistance in *Salmonella* serovars isolated from retail meats in Alberta, Canada. Food Microbiol. 2012 Oct 1; 32(1):110–7. https://doi.org/10.1016/j.fm.2012.04.017

Bäumler AJ, Tsolis RM, Heffron F. The lpf fimbrial operon mediates adhesion of *Salmonella* typhimurium to murine Peyer’s patches. Proc Natl Acad Sci. 1996 Jan 9; 93(1):279–83. https://doi.org/10.1073/pnas.93.1.279

Bäumler AJ, Heffron F. Identification and sequence analysis of lpfABCDE, a putative fimbrial operon of *Salmonella* typhimurium. J Bacteriol. 1995 Apr 1; 177(8):2087–97. https://dx.doi.org/10.1128%2Fjb.177.8.2087-2097.1995

Batchelor M, Hopkins K, Threlfall EJ, Clifton-Hadley FA, Stallwood AD, Davies RH, et al. blaCTX-M genes in clinical *Salmonella* isolates recovered from humans in England and Wales from 1992 to 2003. Antimicrob Agents Chemother. 2005 Apr 1; 49(4):1319–22. https://dx.doi.org/10.1128%2FAAC.49.4.1319-1322.2005

Behravesh CB, Brinson D, Hopkins BA, Gomez TM. Backyard poultry flocks and salmonellosis: a recurring, yet preventable public health challenge. Clin Infect Dis. 2014 May 15; 58(10):1432–8. https://doi.org/10.1093/cid/ciu067

Biswas S, Li Y, Elbediwi M, Yue M. Emergence and dissemination of mcr-carrying clinically relevant *Salmonella* Typhimurium monophasic clone ST34. Microorganisms. 2019 Sep; 7(9):298. https://doi.org/10.3390/microorganisms7090298

Borges KA, Furian TQ, Borsoi A, Moraes HL, Salle CT, Nascimento VP. Detection of virulence-associated genes in *Salmonella* Enteritidis isolates from chicken in South of Brazil. Pesqui Vet Bras. 2013 Dec; 33(12):1416–22.

Borsoi A, Santin E, Santos LR, Salle CT, Moraes HL, Nascimento VP. Inoculation of newly hatched broiler chicks with two Brazilian isolates of *Salmonella* Heidelberg strains with different virulence gene profiles, antimicrobial resistance, and pulsed field gel electrophoresis patterns to intestinal changes evaluation. Poult Sci. 2009 Apr 1; 88(4):750–8. https://doi.org/10.3382/ps.2008-00466

Bupasha ZB, Begum R, Karmakar S, Akter R, Bayzid M, Ahad A, et al.. Multidrug-Resistant *Salmonella* spp. Isolated from Apparently Healthy Pigeons in a Live Bird Market in Chattogram, Bangladesh. World’s Vet J. 2020 Dec 25; 10(4):508–13. https://dx.doi.org/10.29252/scil.2020.wvj61

Campioni F, Bergamini AM, Falcão JP. Genetic diversity, virulence genes and antimicrobial resistance of *Salmonella* Enteritidis isolated from food and humans over a 24-year period in Brazil. Food Microbiol. 2012 Dec 1; 32(2):254–64. https://doi.org/10.1016/j.fm.2012.06.008

Cardona-Castro N, Restrepo-Pineda E, Correa-Ochoa M. Detection of hilA gene sequences in serovars of *Salmonella* enterica sufigbspecies enterica. Mem Inst Oswaldo Cruz. 2002 Dec; 97(8):1153–6. https://doi.org/10.1590/s0074-02762002000800016

Castilla KS, Ferreira CS, Moreno AM, Nunes IA, Ferreira AJ. Distribution of virulence genes sefC, pefA and spvC in *Salmonella* Enteritidis phage type 4 strains isolated in Brazil. Braz J Microbiol. 2006 Jun; 37(2):135–9. https://doi.org/10.1590/S1517-83822006000200007

Cesco MAO, Zimermann FC, Giotto DB, Guayba J, Borsoi A, Rocha SLS, et al. Pesquisa de Genes de Virulência em Salmonella Hadar em Amostras Provenientes de Material Avícola; R0701-0; Anais 35 Congresso Brasileiro de Medicina Veterinária: Porto Alegre, Brazil, 2008.

Clinical and Laboratory Standards Institute (CLSI). Performance Standards for Antimicrobial Susceptibility Testing; Approved Standard, 28th ed.; Document M100, 2019; Clinical and Laboratory Standards Institute: Wayne, PA, USA, 2019.

Collinson SK, Doig PC, Doran JL, Clouthier S, Kay WW. Thin, aggregative fimbriae mediate binding of *Salmonella* enteritidis to fibronectin. J Bacteriol. 1993 Jan 1; 175(1):12–8. https://dx.doi.org/10.1128%2Fjb.175.1.12-18.1993

Crăciunaş C, Keul AL, Flonta M, Cristea M. DNA-based diagnostic tests for *Salmonella* strains targeting hilA, agfA, spvC and sef genes. J Environ Manag. 2012 Mar 1; 95: S15–8. https://doi.org/10.1016/j.jenvman.2010.07.027

Dallenne C, Da Costa A, Decré D, Favier C, Arlet G. Development of a set of multiplex PCR assays for the detection of genes encoding important β-lactamases in Enterobacteriaceae. J. Antimicrob. Chemother. 2010 Mar 1; 65(3):490–5. https://doi.org/10.1093/jac/dkp498

Daniel WW. Biostatistics: A Foundation for Analysis in the Health Sciences, 7th ed.; John Willey & Sons: New York, NY, USA; 1999.

Darwin KH, Miller VL. InvF is required for expression of genes encoding proteins secreted by the SPI1 type III secretion apparatus in *Salmonella* typhimurium. J Bacteriol. 1999 Aug 15; 181(16):4949–54. https://doi.org/10.1128/jb.181.16.4949-4954.1999

de Freitas CG, Santana ÂP, da Silva PH, Gonçalves VS, Barros MD, Torres FA, et al. PCR multiplex for detection of *Salmonella* Enteritidis, Typhi and Typhimurium and occurrence in poultry meat. Int J Food Microbiol. 2010 Apr 30; 139(1-2):15–22. https://doi.org/10.1016/j.ijfoodmicro.2010.02.007

Doosti A, Mahmoudi E, Jami MS, Mokhtari-Farsani A. Prevalence of aadA1, aadA2, aadB, strA and strB genes and their associations with multidrug resistance phenotype in *Salmonella* Typhimurium isolated from poultry carcasses. Thai J Vet Med. 2016; 46(4):691–7. https://he01.tci-thaijo.org/index.php/tjvm/article/view/73827

Ed-Dra A, Karraouan B, El Allaoui A, Khayatti M, El Ossmani H, Filali FR, et al. Antimicrobial resistance and genetic diversity of *Salmonella* Infantis isolated from foods and human samples in Morocco. J Glob Antimicrob Resist. 2018 Sep 1; 14:297–301. https://doi.org/10.1016/j.jgar.2018.05.019

Elbediwi M, Pan H, Biswas S, Li Y, Yue M. Emerging colistin resistance in *Salmonella* enterica serovar Newport isolates from human infections. Emerg Microbes Infect. 2020 Jan 1; 9(1):535–8. https://dx.doi.org/10.1080%2F22221751.2020.1733439

Elkenany R, Elsayed MM, Zakaria AI, El-Sayed SA, Rizk MA. Antimicrobial resistance profiles and virulence genotyping of *Salmonella* enterica serovars recovered from broiler chickens and chicken carcasses in Egypt. BMC Vet Res. 2019 Dec; 15(1):1–9. https://doi.org/10.1186/s12917-019-1867-z

El-Sharkawy H, Tahoun A, El-Gohary AE, El-Abasy M, El-Khayat F, Gillespie T, et al. Epidemiological, molecular characterization and antibiotic resistance of *Salmonella* enterica serovars isolated from chicken farms in Egypt. Gut Pathog. 2017 Dec; 9(1):1–2. https://doi.org/10.1186/s13099-017-0157-1

Garedew L, Hagos Z, Addis Z, Tesfaye R, Zegeye B. Prevalence and antimicrobial susceptibility patterns of *Salmonella* isolates in association with hygienic status from butcher shops in Gondar town, Ethiopia. Antimicrob Resist Infect Control. 2015 Dec; 4(1):1–7. https://doi.org/10.1186/s13756-015-0062-7

Gieraltowski L, Higa J, Peralta V, Green A, Schwensohn C, Rosen H, et al. National outbreak of multidrug resistant *Salmonella* Heidelberg infections linked to a single poultry company. PloS one. 2016 Sep 15; 11(9):e0162369. https://doi.org/10.1371/journal.pone.0162369

González F, Araque M. Association of transferable quinolone resistance determinant qnrB19 with extended-spectrum β-lactamases in *Salmonella* Give and Salmonella Heidelberg in Venezuela. Int J Microbiol. 2013 Oct; 2013. https://doi.org/10.1155/2013/628185

Guo X, Chen J, Beuchat LR, Brackett RE. PCR detection of *Salmonella* enterica serotype Montevideo in and on raw tomatoes using primers derived from hila. Appl Environ Microbiol. 2000 Dec 1; 66(12):5248–52. https://dx.doi.org/10.1128%2Faem.66.12.5248-5252.2000

Hawker J, Begg N, Reintjes R, Ekdahl K, Edeghere O, Van Steenbergen JE. Communicable disease control and health protection handbook. John Wiley & Sons; 2018 Dec 3. https://doi.org/10.1002/9781119328070.ch2

Heidary M, Momtaz H, Madani M. Characterization of diarrheagenic antimicrobial resistant *Escherichia coli* isolated from pediatric patients in Tehran, Iran. Iran. Red. Crescent. Med. J. 2014 Apr; 16 (4)

Huehn S, La Ragione RM, Anjum M, Saunders M, Woodward MJ, Bunge C, et al. Virulotyping and antimicrobial resistance typing of *Salmonella* enterica serovars relevant to human health in Europe. Foodborne Pathog Dis. 2010 May 1; 7(5):523–35. https://doi.org/10.1089/fpd.2009.0447

Interagency Coordination Group (IACG). No Time to Wait: Securing the Future from Drug-Resistant Infections, Report to the Secretary General of the United Nations; Interagency Coordination Group: New York, NY, USA; 2019.

ISO, 6579: 2002. Microbiology of Food and Animal Feeding Stuffs. Horizontal Method for the Detection of Salmonella spp.; British Standard Institute: London, UK; 2002.

Im MC, Jeong SJ, Kwon YK, Jeong OM, Kang MS, Lee YJ. Prevalence and characteristics of *Salmonella* spp. isolated from commercial layer farms in Korea. Poult Sci. 2015 Jul 1; 94(7):1691–8. https://doi.org/10.3382/ps/pev137

Issenhuth-Jeanjean S, Roggentin P, Mikoleit M, Guibourdenche M, de Pinna E, Nair S, et al. Supplement 2008–2010 (no. 48) to the white–Kauffmann–Le minor scheme. Res Microbiol. 2014 Sep 1; 165(7):526–30. https://doi.org/10.1016/j.resmic.2014.07.004

Jahantigh M, Samadi K, Dizaji RE, Salari S. Antimicrobial resistance and prevalence of tetracycline resistance genes in *Escherichia coli* isolated from lesions of colibacillosis in broiler chickens in Sistan, Iran. BMC Vet Res. 2020 Dec; 16(1):1–6. https://doi.org/10.1186/s12917-020-02488-z

Jain P, Chowdhury G, Samajpati S, Basak S, Ganai A, Samanta S, et al. Characterization of non-typhoidal *Salmonella* isolates from children with acute gastroenteritis, Kolkata, India, during 2000–2016. Braz J Microbiol. 2020 Jun; 51(2):613–27. https://doi.org/10.1007/s42770-019-00213-z

Jaja IF, Bhembe NL, Green E, Oguttu J, Muchenje V. Molecular characterisation of antibiotic-resistant *Salmonella* enterica isolates recovered from meat in South Africa. Acta trop. 2019 Feb 1; 190:129–36. https://doi.org/10.1016/j.actatropica.2018.11.003

Jiang Z, Paudyal N, Xu Y, Deng T, Li F, Pan H, et al. Antibiotic resistance profiles of *Salmonella* recovered from finishing pigs and slaughter facilities in Henan, China. Front Microbiol. 2019 Jul 4; 10:1513. https://doi.org/10.3389/fmicb.2019.01513

Karim MR, Giasuddin M, Samad MA, Mahmud MS, Islam MR, Rahman MH, et al. Prevalence of *Salmonella* spp. in poultry and poultry products in Dhaka, Bangladesh. Int J Anim Biol. 2017; 3(4):18–22.

Kaushik P, Kumari S, Bharti SK, Dayal S. Isolation and prevalence of *Salmonella* from chicken meat and cattle milk collected from local markets of Patna, India. Vet World. 2014 Feb 1; 7(2):62. https://dx.doi.org/10.14202/vetworld.2014.62-65

Keerthirathne TP, Ross K, Fallowfield H, Whiley H. Reducing risk of salmonellosis through egg decontamination processes. Int J Environ Res Public Health. 2017 Mar; 14(3):335. https://doi.org/10.3390/ijerph14030335

Kingsley RA, Humphries AD, Weening EH, De Zoete MR, Winter S, Papaconstantinopoulou A, et al. Molecular and phenotypic analysis of the CS54 island of *Salmonella* enterica serotype Typhimurium: identification of intestinal colonization and persistence determinants. Infect Immun. 2003 Feb 1; 71(2):629–40. https://doi.org/10.1128/iai.71.2.629-640.2003

Kirk MD, Pires SM, Black RE, Caipo M, Crump JA, Devleesschauwer B, et al. World Health Organization estimates of the global and regional disease burden of 22 foodborne bacterial, protozoal, and viral diseases, 2010: a data synthesis. PLoS Med. 2015 Dec 3; 12(12):e1001921. https://doi.org/10.1371/journal.pmed.1001921

Kozak GK, Boerlin P, Janecko N, Reid-Smith RJ, Jardine C. Antimicrobial resistance in *Escherichia coli* isolates from swine and wild small mammals in the proximity of swine farms and in natural environments in Ontario, Canada. Appl Environ Microbiol. 2009 Feb 1; 75(3):559–66. https://doi.org/10.1128/aem.01821-08

Lin D, Yan M, Lin S, Chen S. Increasing prevalence of hydrogen sulfide negative *Salmonella* in retail meats. Food Microbiol. 2014 Oct 1; 43:1–4. https://doi.org/10.1016/j.fm.2014.04.010

Long M, Lai H, Deng W, Zhou K, Li B, Liu S, et al. Disinfectant susceptibility of different Sal*m*onella serotypes isolated from chicken and egg production chains. J Appl Microbiol. 2016 Sep; 121(3):672–81. https://doi.org/10.1111/jam.13184

Lu Y, Wu CM, Wu GJ, Zhao HY, He T, Cao XY, et al. Prevalence of antimicrobial resistance among *Salmonella* isolates from chicken in China. Foodborne Pathog Dis. 2011 Jan 1; 8(1):45–53. https://doi.org/10.1089/fpd.2010.0605

Mahmud MS, Bari ML, Hossain MA. Prevalence of Salmonella serovars and antimicrobial resistance profiles in poultry of Savar area, Bangladesh. Foodborne Pathog Dis. 2011 Oct 1; 8(10):1111–8. https://doi.org/10.1089/fpd.2011.0917

Magiorakos AP, Srinivasan A, Carey RT, Carmeli Y, Falagas MT, Giske CT, et al. Multidrug-resistant, extensively drug-resistant and pandrug-resistant bacteria: an international expert proposal for interim standard definitions for acquired resistance. Clin Microbiol Infect. 2012 Mar 1; 18(3):268–81. https://doi.org/10.1111/j.1469-0691.2011.03570.x

Malorny B, Hoorfar J, Hugas M, Heuvelink A, Fach P, Ellerbroek L, et al. Interlaboratory diagnostic accuracy of a *Salmonella* specific PCR-based method. Int J Food Microbiol. 2003 Dec 31; 89(2-3):241–9. https://doi.org/10.1016/s0168-1605(03)00154-5

Manyi-Loh C, Mamphweli S, Meyer E, Okoh A. Antibiotic use in agriculture and its consequential resistance in environmental sources: potential public health implications. Molecules. 2018 Apr; 23(4):795. https://doi.org/10.3390/molecules23040795

McCrea BA, Tonooka KH, VanWorth C, Atwill ER, Schrader JS, Boggs CL. Prevalence of *Campylobacter* and *Salmonella* species on farm, after transport, and at processing in specialty market poultry. Poult Sci. 2006 Jan 1; 85(1):136–43. https://doi.org/10.1093/ps/85.1.136

Momtaz S, Saha O, Usha MK, Sultana M, Hossain MA. Occurrence of pathogenic and multidrug resistant *Salmonella* spp. in poultry slaughter-house in Bangladesh. Bioresearch Communications- (BRC). 2018 Jul 1; 4(2):506–15.

Motta RN, Oliveira MM, Magalhães PS, Dias AM, Aragao LP, Forti AC, et al. Plasmid-mediated extended-spectrum beta-lactamase-producing strains of Enterobacteriaceae isolated from diabetes foot infections in a Brazilian diabetic center. Braz J Infect Dis. 2003 Apr; 7(2):129–34. https://doi.org/10.1590/s1413-86702003000200006

Mridha D, Uddin MN, Alam B, Akhter AT, Islam SS, Islam MS, et al. Identification and characterization of *Salmonella* spp. from samples of broiler farms in selected districts of Bangladesh. Vet World. 2020 Feb; 13(2):275. https://doi.org/10.14202/vetworld.2020.275-283

Naing L, Winn T, Rusli BN. Practical issues in calculating the sample size for prevalence studies. Arch Orofac Sci. 2006; 1:9–14. https://www.scribd.com/doc/63105077/How-to-Calculate-Sample-Size

Nayak R, Stewart T, Wang RF, Lin J, Cerniglia CE, Kenney PB. Genetic diversity and virulence gene determinants of antibiotic-resistant *Salmonella* isolated from preharvest turkey production sources. Int J Food Microbiol. 2004 Feb 15; 91(1):51–62. https://doi.org/10.1016/s0168-1605(03)00330-1

Nidaullah H, Abirami N, Shamila-Syuhada AK, Chuah LO, Nurul H, Tan TP, et al. Prevalence of *Salmonella* in poultry processing environments in wet markets in Penang and Perlis, Malaysia. Vet World. 2017 Mar; 10(3):286. https://doi.org/10.14202/vetworld.2017.286-292

O’Bryan CA, Crandall PG, Ricke SC. Antimicrobial resistance in foodborne pathogens. Food and Feed Safety Systems and Analysis. 2018 Jan 1; 99-115. https://doi.org/10.1016/C2016-0-00136-5

O’Neill J. Review on Antimicrobial Resistance Antimicrobial Resistance: Tackling a crisis for the health and wealth of nations. London: Review on Antimicrobial Resistance. 2014. Available from: https://amr-review.org/sites/default/files/AMR%20Review%20Paper%20%20Tackling%20a%20crisis%20for%20the%20health%20and%20wealth%20of%20nations_1.pdf

Oliveira SD, Santos LR, Schuch DM, Silva AB, Salle CT, Canal CW. Detection and identification of salmonellas from poultry-related samples by PCR. Vet Microbiol. 2002 Jun 5; 87(1):25–35. https://doi.org/10.1016/s0378-1135(02)00028-7

Osimani A, Aquilanti L, Clementi F. Salmonellosis associated with mass catering: a survey of European Union cases over a 15-year period. Epidemiol Infect. 2016 Oct; 144(14):3000–12. https://doi.org/10.1017/s0950268816001540

Pang L, Zhang Z, Xu J. Surveillance of foodborne disease outbreaks in China in 2006—2010. Chin J Food Hyg. 2011; 23:560–3.

Park HC, Baig IA, Lee SC, Moon JY, Yoon MY. Development of ssDNA aptamers for the sensitive detection of *Salmonella* typhimurium and *Salmonella* enteritidis. Appl Biochem Biotechnol. 2014 Sep; 174(2):793–802. https://doi.org/10.1007/s12010-014-1103-z

Parvej MS, Rahman M, Uddin MF, Nazir KN, Jowel MS, Khan MF, et al. Isolation and characterization of *Salmonella* enterica serovar typhimurium circulating among healthy chickens of Bangladesh. Turkish Journal of Agriculture-Food Science and Technology. 2016 Jul 15; 4(7):519–23. https://doi.org/10.24925/turjaf.v4i7.519-523.695

Parvej MS, Nazir KN, Rahman MB, Jahan M, Khan MF, Rahman M. Prevalence and characterization of multi-drug resistant Salmonella Enterica serovar Gallinarum biovar Pullorum and Gallinarum from chicken. Vet World. 2016 Jan; 9(1):65. https://doi.org/10.14202/vetworld.2016.65-70

Parvin MS, Hasan MM, Ali MY, Chowdhury EH, Rahman MT, Islam MT. Prevalence and Multidrug Resistance Pattern of *Salmonella* Carrying Extended-Spectrum β-Lactamase in Frozen Chicken Meat in Bangladesh. J Food Prot. 2020 Dec; 83(12):2107–21. https://doi.org/10.4315/jfp-20-172

Paterson DL, Bonomo RA. Extended-spectrum β-lactamases: a clinical update. Clin Microbiol Rev. 2005 Oct 1; 18(4):657–86. https://doi.org/10.1128/cmr.18.4.657-686.2005

Paudyal N, Pan H, Wu B, Zhou X, Zhou X, Chai W, Wu Q, Li S, Li F, Gu G, Wang H. Persistent asymptomatic human infections by Salmonella enterica serovar Newport in China. Msphere. 2020 Jun 24; 5(3). https://doi.org/10.1128/msphere.00163-20

Prager R, Rabsch W, Streckel W, Voigt W, Tietze E, Tschäpe H. Molecular properties of Salmonella enterica serotype Paratyphi B distinguish between its systemic and its enteric pathovars. J Clin Microbiol. 2003 Sep 1; 41(9):4270–8. https://doi.org/10.1128/jcm.41.9.4270-4278.2003

Ramatla T, Ngoma L, Adetunji M, Mwanza M. Evaluation of antibiotic residues in raw meat using different analytical methods. Antibiotics. 2017 Dec; 6(4):34. https://doi.org/10.3390/antibiotics6040034

Robicsek A, Strahilevitz J, Sahm DF, Jacoby GA, Hooper DC. qnr prevalence in ceftazidime-resistant Enterobacteriaceae isolates from the United States. Antimicrob Agents Chemother. 2006 Aug 1; 50(8):2872–4. https://doi.org/10.1128/aac.01647-05

Rusul G, Khair J, Radu S, Cheah CT, Yassin RM. Prevalence of Salmonella in broilers at retail outlets, processing plants and farms in Malaysia. Int J Food Microbiol. 1996 Dec 1; 33(2-3):183–94. https://doi.org/10.1016/0168-1605(96)01125-7

Saitanu K, Jerngklinchan J, Koowatananukul C. Incidence of salmonellae in duck eggs in Thailand. Southeast Asian J Trop Med Public Health. 1994 Jun 1; 25:328–31. PMID: 7855651

Sallam KI, Mohammed MA, Hassan MA, Tamura T. Prevalence, molecular identification and antimicrobial resistance profile of *Salmonella* serovars isolated from retail beef products in Mansoura, Egypt. Food control. 2014 Apr 1; 38:209–14. https://doi.org/10.1016/j.foodcont.2013.10.027

Samanta I, Joardar SN, Das PK, Das P, Sar TK, Dutta TK, et al. Virulence repertoire, characterization, and antibiotic resistance pattern analysis of *Escherichia coli* isolated from backyard layers and their environment in India. Avian Dis. 2014 Mar;58(1):39–45. https://doi.org/10.1637/10586-052913-reg.1

Sarker BR, Ghosh S, Chowdhury S, Dutta A, Chandra Deb L, Krishna Sarker B, et al. Prevalence and antimicrobial susceptibility profiles of non-typhoidal *Salmonella* isolated from chickens in Rajshahi, Bangladesh. Vet Med Sci. 2021 May; 7(3):820–830. https://doi.org/10.1002/vms3.440

Sarker MS, Ahad A, Ghosh SK, Mannan MS, Sen A, Islam S, et al. Antibiotic-resistant *Escherichia coli* in deer and nearby water sources at safari parks in Bangladesh. Vet World. 2019 Oct; 12(10):1578. https://doi.org/10.14202/vetworld.2019.1578-1583

Shah AH, Korejo NA. Antimicrobial resistance profile of *Salmonella* serovars isolated from chicken meat. J Vet Anim Sci. 2012; 2:40–6.

Sharma J, Kumar D, Hussain S, Pathak A, Shukla M, Kumar VP, et al. Prevalence, antimicrobial resistance and virulence genes characterization of nontyphoidal *Salmonella* isolated from retail chicken meat shops in Northern India. Food control. 2019 Aug 1; 102:104–11. http://dx.doi.org/10.1016/j.foodcont.2019.01.021

Siddiky NA, Sarker MS, Khan M, Rahman S, Begum R, Kabir M, et al. Virulence and Antimicrobial Resistance Profiles of *Salmonella* enterica Serovars Isolated from Chicken at Wet Markets in Dhaka, Bangladesh. Microorganisms. 2021 May; 9(5):952. https://doi.org/10.3390/microorganisms9050952

Silva C, Puente JL, Calva E. *Salmonella* virulence plasmid: pathogenesis and ecology. Pathog Dis. 2017 Aug;75(6):ftx070. https://doi.org/10.1093/femspd/ftx070

Sin M, Yoon S, Kim YB, Noh EB, Seo KW, Lee YJ. Molecular characteristics of antimicrobial resistance determinants and integrons in *Salmonella* isolated from chicken meat in Korea. J Appl Poult Res. 2020 Jun 1; 29(2):502–14. https://doi.org/10.1016/j.japr.2019.12.010

Sobur A, Hasan M, Haque E, Mridul AI, Noreddin A, El Zowalaty ME, et al. Molecular detection and antibiotyping of multidrug-resistant *Salmonella* isolated from houseflies in a fish market. Pathogens. 2019 Dec; 8(4):191. https://doi.org/10.3390/pathogens8040191

Spector MP, Kenyon WJ. Resistance and survival strategies of Salmonella enterica to environmental stresses. Food Res Int. 2012 Mar 1; 45(2):455–81. https://doi.org/10.1016/j.foodres.2011.06.056

Sripaurya B, Ngasaman R, Benjakul S, Vongkamjan K. Virulence genes and antibiotic resistance of *Salmonella* recovered from a wet market in Thailand. J Food Saf. 2019 Apr; 39(2):e12601. https://doi.org/10.1111/jfs.12601

Sultana M, Bilkis R, Diba F, Hossain MA. Predominance of multidrug resistant zoonotic Salmonella Enteritidis genotypes in poultry of Bangladesh. J Poult Sci. 2014:0130222. https://doi.org/10.2141/jpsa.0130222

Suresh Y, Kiranmayi CB, Rao TS, Srivani M, Subhashini N, Chaitanya G, et al. Multidrug resistance and ESBL profile of *Salmonella* serovars isolated from poultry birds and foods of animal origin. The Pharma Innovation Journal. 2019; 8(4):277–82

Suresh T, Hatha AA, Sreenivasan D, Sangeetha N, Lashmanaperumalsamy P. Prevalence and antimicrobial resistance of *Salmonella* enteritidis and other salmonellas in the eggs and egg-storing trays from retails markets of Coimbatore, South India. Food Microbiol. 2006 May 1; 23(3):294–9. https://doi.org/10.1016/j.fm.2005.04.001

Swamy SC, Barnhart HM, Lee MD, Dreesen DW. Virulence determinants *inv*A and *spv*C in salmonellae isolated from poultry products, wastewater, and human sources. Appl Environ Microbiol. 1996 Oct 1; 62(10):3768–71. https://doi.org/10.1128/aem.62.10.3768-3771.1996

Takaya A, Yamamoto T, Tokoyoda K. Humoral immunity vs. *Salmonella*. Front Immunol. 2020 Jan 21; 10:3155. https://doi.org/10.3389/fimmu.2019.03155

Tarabees R, Elsayed MS, Shawish R, Basiouni S, Shehata AA. Isolation and characterization of *Salmonella* Enteritidis and *Salmonella* Typhimurium from chicken meat in Egypt. J Infect Dev Ctries. 2017 Apr 30; 11(04):314–9. https://doi.org/10.3855/jidc.8043

Thung TY, Radu S, Mahyudin NA, Rukayadi Y, Zakaria Z, Mazlan N, et al. Prevalence, virulence genes and antimicrobial resistance profiles of *Salmonella* serovars from retail beef in Selangor, Malaysia. Front Microbiol. 2018 Jan 11; 8:2697. https://doi.org/10.3389/fmicb.2017.02697

Titilawo Y, Sibanda T, Obi L, Okoh A. Multiple antibiotic resistance indexing of *Escherichia coli* to identify high-risk sources of faecal contamination of water. Environ Sci Pollut Res Int. 2015 Jul; 22(14):10969–80. https://doi.org/10.1007/s11356-014-3887-3

Van TT, Chin J, Chapman T, Tran LT, Coloe PJ. Safety of raw meat and shellfish in Vietnam: an analysis of *Escherichia coli* isolations for antibiotic resistance and virulence genes. Int J Food Microbiol. 2008 Jun 10; 124(3):217–23. https://doi.org/10.1016/j.ijfoodmicro.2008.03.029

Vuthy Y, Lay KS, Seiha H, Kerleguer A, Aidara-Kane A. Antibiotic susceptibility and molecular characterization of resistance genes among *Escherichia coli* and among *Salmonella* subsp. in chicken food chains. Asian Pac J Trop Biomed. 2017 Jul 1; 7(7):670–4. https://doi.org/10.1016/j.apjtb.2017.07.002

Waghamare RN, Paturkar AM, Zende RJ, Vaidya VM, Gandage RS, Aswar NB, et al. Studies on occurrence of invasive *Salmonella* spp. from unorganised poultry farm to retail chicken meat shops in Mumbai city, India. Int J Curr Microbiol Appl Sci. 2017; 6(5):630–41. https://doi.org/10.20546/ijcmas.2017.605.073

Wang H, Ye K, Wei X, Cao J, Xu X, Zhou G. Occurrence, antimicrobial resistance and biofilm formation of *Salmonella* isolates from a chicken slaughter plant in China. Food Control. 2013 October; 33:378–84. https://doi.org/10.1016/j.foodcont.2013.03.030

World Health Organization (WHO). WHO Releases the 2019 AWaRe Classification Antibiotics; World Health Organization: New York, NY, USA; 2019.

Xiang Y, Li F, Dong N, Tian S, Zhang H, Du X, et al. Investigation of a Salmonellosis outbreak caused by multidrug resistant *Salmonella* Typhimurium in China. Front Microbiol. 2020 Apr 29; 11:801. https://doi.org/10.3389/fmicb.2020.00801

Xu Y, Zhou X, Jiang Z, Qi Y, Ed-Dra A, Yue M. Epidemiological investigation and antimicrobial resistance profiles of *Salmonella* isolated from breeder chicken hatcheries in Henan, China. Front Cell Infect Microbiol. 2020 Sep 15; 10:497. https://doi.org/10.3389/fcimb.2020.00497

Yang B, Qu D, Zhang X, Shen J, Cui S, Shi Y, et al. Prevalence and characterization of *Salmonella* serovars in retail meats of marketplace in Shaanxi, China. Int J Food Microbiol. 2010 Jun 30; 141(1-2):63–72. https://doi.org/10.1016/j.ijfoodmicro.2010.04.015

Yoo AY, Yu JE, Yoo H, Lee TH, Lee WH, Oh JI, et al. Role of sigma factor E in regulation of *Salmonella* Agf expression. Biochem Biophys Res Commun. 2013 Jan 4; 430(1):131–6. https://doi.org/10.1016/j.bbrc.2012.11.025

Yu H, Elbediwi M, Zhou X, Shuai H, Lou X, Wang H, et al. Epidemiological and genomic characterization of *Campylobacter jejuni* isolates from a foodborne outbreak at Hangzhou, China. Int J Mol Sci. 2020 Jan; 21(8):3001. https://doi.org/10.3390/ijms21083001

Zahraei SM, Mahzoniae MR, Ashrafi A. Amplification of invA gene of *Salmonalla* by polymerase chain reaction (PCR) as a specific method for detection of salmonellae. J Vet Res. 61(2):195–199. https://www.sid.ir/en/journal/ViewPaper.aspx?id=83064

Zeng YB, Xiong LG, Tan MF, Li HQ, Yan H, Zhang L, et al. Prevalence and antimicrobial resistance of *Salmonella* in pork, chicken, and duck from retail markets of China. Foodborne Pathog Dis. 2019 May 1; 16(5):339–45. https://doi.org/10.1089/fpd.2018.2510

Zhu Y, Lai H, Zou L, Yin S, Wang C, Han X, et al. Antimicrobial resistance and resistance genes in *Salmonella* strains isolated from broiler chickens along the slaughtering process in China. Int J Food Microbiol. 2017 Oct 16; 259:43–51. https://doi.org/10.1016/j.ijfoodmicro.2017.07.023

